# An overcomplete approach to fitting drift-diffusion decision models to trial-by-trial data

**DOI:** 10.1101/2020.01.30.925123

**Authors:** Q. Feltgen, J. Daunizeau

## Abstract

Drift-diffusion models or DDMs are becoming a standard in the field of computational neuroscience. They extend models from signal detection theory by proposing a simple mechanistic explanation for the observed relationship between decision outcomes and reaction times (RT). In brief, they assume that decisions are triggered once the accumulated evidence in favor of a particular alternative option has reached a predefined threshold. Fitting a DDM to empirical data then allows one to interpret observed group or condition differences in terms of a change in the underlying model parameters. However, current approaches only yield reliable parameter estimates in specific situations (c.f. fixed drift rates vs drift rates varying over trials). In addition, they become computationally unfeasible when more general DDM variants are considered (e.g., with collapsing bounds). In this note, we propose a fast and efficient approach to parameter estimation that relies on fitting a “self-consistency” equation that RT fulfill under the DDM. This effectively bypasses the computational bottleneck of standard DDM parameter estimation approaches, at the cost of estimating the trial-specific neural noise variables that perturb the underlying evidence accumulation process. For the purpose of behavioral data analysis, these act as nuisance variables and render the model “overcomplete”, which is finessed using a variational Bayesian system identification scheme. But for the purpose of neural data analysis, estimates of neural noise perturbation terms are a desirable (and unique) feature of the approach. Using numerical simulations, we show that this “overcomplete” approach matches the performance of current parameter estimation approaches for simple DDM variants, and outperforms them for more complex DDM variants. Finally, we demonstrate the added-value of the approach, when applied to a recent value-based decision making experiment.

## 1. Introduction

Over the past two decades, neurocognitive processes of decision making have been extensively studied under the framework of so-called *drift-diffusion models* or DDMs. These models tie together decision outcomes and response times (RT) by assuming that decisions are triggered once the accumulated evidence in favor of a particular alternative option has reached a predefined threshold (Ratcliff and McKoon, 2008; Ratcliff et al., 2016). They owe their popularity both to experimental successes in capturing observed data in a broad set of behavioral studies (De Martino et al., 2012; Gold and Shadlen, 2007; Hanks et al., 2014; Milosavljevic et al., 2010; Pedersen et al., 2017; Resulaj et al., 2009), and to theoretical work showing that DDMs can be thought of as optimal decision problem solvers (Balci et al., 2011; Bogacz et al., 2006; Drugowitsch et al., 2012; Tajima et al., 2016; Zhang, 2012). Critically, mathematical analyses of the DDM soon demonstrated that it suffers from inherent non-identifiability issues, e.g., predicted choices and RTs are invariant under any arbitrary rescaling of DDM parameters (Ratcliff and Tuerlinckx, 2002; Ratcliff et al., 2016). This is important because, in principle, this precludes proper, quantitative, DDM-based data analysis. Nevertheless, over the past decade, many statistical approaches to DDM parameter estimation have been proposed, which yield efficient parameter estimation under simplifying assumptions (Grasman et al., 2009; Hawkins et al., 2015; Pedersen and Frank, 2020; Vandekerckhove and Tuerlinckx, 2008; Voskuilen et al., 2016; Voss and Voss, 2007; Wagenmakers et al., 2007, 2008; Wiecki et al., 2013; Zhang, 2012; Zhang et al., 2014). Typically, these techniques essentially fit the choice-conditional distribution of observed RT (or moments thereof), having arbitrary fixed at least one of the DDM parameters. They are now established statistical tools for experimental designs where the observed RT variability is mostly induced by internal (e.g., neural) stochasticity in the decision process (Boehm et al., 2018).

Now current decision making experiments typically consider situations in which decision-relevant variables are manipulated on a trial-by-trial basis. For example, the reliability of perceptual evidence (e.g., the psychophysical contrast in a perceptual decision) may be systematically varied from one trial to the next. Under current applications of the DDM, this implies that some internal model variables (e.g., the drift rate) effectively vary over trials. Classical DDM parameter estimation approaches do not optimally handle this kind of experimental designs, because these lack the trial repetitions that would be necessary to provide empirical estimates of RT moments in each condition. In turn, alternative statistical approaches to parameter estimation have been proposed, which can exploit predictable inter-trial variations of DDM variables to fit the model to RT data (Fontanesi et al., 2019a, 2019b; Gluth and Meiran, 2019; Moens and Zenon, 2017; Pedersen et al., 2017; Wabersich and Vandekerckhove, 2014). In brief, they directly compare raw RT data with expected RTs, which vary over trials in response to known variations in internal variables. Although close to optimal from a statistical perspective, they suffer from a challenging computational bottleneck that lies in the trial-by-trial derivation of RT first-order moments. This is why they are typically constrained to simple DDM variants, for which analytical solutions exist (Fengler et al., 2020; Navarro and Fuss, 2009; Shinn et al., 2020; Srivastava et al., 2016).

This note is concerned with the issue of obtaining reliable DDM parameter estimates from concurrent trial-by-trial choice and response time data, for a broad class of DDM variants. We propose a fast and efficient approach that relies on fitting a *self-consistency* equation, which RTs necessarily fulfill under the DDM. This provides a simple and elegant way to bypassing the common computational bottleneck of existing approaches, at the cost of considering additional trial-specific nuisance model variables. These are the cumulated “neural” noise that perturbs the evidence accumulation process at the corresponding trial. Including these variables in the model makes it “overcomplete”, the identification of which is finessed with a dedicated variational Bayesian scheme. In turn, the ensuing overcomplete approach generalizes to a large class of DDM model variants, without any additional computational and/or implementational burden.

In section 2 of this document, we briefly recall the derivation of the DDM, and summarize the impact of DDM parameters onto the conditional RT distributions. In sections 3 and 4, we derive the DDM's self-consistency equation and describe the ensuing overcomplete approach to DDM-based data analysis. In section 5, we perform parameter recovery analyses for standard DDM fitting procedures and the overcomplete approach. In section 6, we demonstrate the added-value of the overcomplete approach, when applied to a value-based decision making experiment. Finally, in section 7, we discuss our results in the context of the existing literature. In particular, we comment on the potential utility of neural noise perturbation estimates for concurrent neuroimaging data analysis.

## 2. Model formulation and impact of DDM parameters

First, let us recall the simplest form of a drift-diffusion decision model or DDM (in what follows, we will refer to this variant as the “vanilla” DDM). Let *x* (*t*) be a decision variable that captures the accumulated evidence (up to time *t*) in favour of a given option in a binary choice set. Under the vanilla DDM, a decision is triggered whenever *x* (*t*) hits either of two bounds, which are positioned at *x* = *b* and *x* = −*b*, respectively. When a bound hit occurs defines the decision time, and which bound is hit determines the (binary) decision outcome *o*. By assumption, the decision variable *x* (*t*) is supposed to follow the following stochastic differential equation:

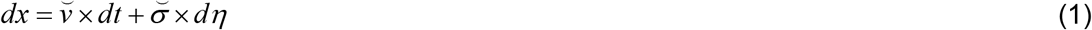

where *v* is the drift rate, *dη* ~ *N*(0, *dt*) is a standard Wiener process, and 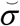 is the standard deviation of the stochastic (diffusion) perturbation term.

Equation 1 can be discretized using an Euler-Maruyma scheme (Kloeden and Platen, 1992), yielding the following discrete form of the decision variable dynamics:

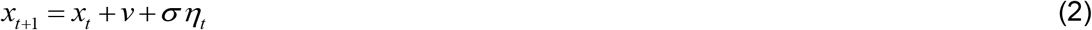

where *t* indexes time on a temporal grid with resolution 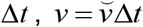 is the discrete-time drift rate, 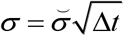 is the discrete-time standard deviation of the perturbation term and *η_t_* ~ *N*(0, 1) is a standard normal random variable. By convention, the system’s initial condition is denoted as *x*_0_, which we refer to as the “initial condition”.

The joint distribution of response times and decision outcomes depend upon the DDM parameters, which include: the drift rate *v*, the bound’s height *b*, the noise’s standard deviation *σ* and the initial condition *x*_0_. DDMs also typically include a so-called “non-decision” time parameter *T*_*ND*_, which captures systematic latencies between covert bound hit times and overt response times. Under such simple DDM variant, variability in response times and decision outcomes derive from stochastic terms *η*. These are typically thought of as neural noise that perturb the evidence accumulation process within the brain’s decision system (Gold and Shadlen, 2007; Guevara Erra et al., 2019; Turner et al., 2015).

Under such simple DDM variant, analytical expressions exist for the first two moments of RT distributions (Srivastava et al., 2016). Higher-order moments can also be derived from efficient semi-analytical solutions to the issue of deriving the joint choice/RT distribution (Navarro and Fuss, 2009). However, more complex variants of the DDM (including, e.g., collapsing bounds) are much more difficult to simulate, and require either sampling schemes or numerical solvers of the underlying Fokker-Planck equation (Fengler et al., 2020; Shinn et al., 2020).

Figures 1 to 4 below demonstrate the impact of model parameters on the decision outcome ratios *P* (*o* | *v*, *x*_0_, *b*,*σ*) and the first three moments of conditional hitting time (HT) distributions, namely: their mean *E* [*HT* | *o*, *v*, *x*_0_, *b*, *σ*, variance *V* [*HT* | *o*, *v*, *x*_0_, *b*,*σ*] and skewness *S_k_* [*HT* | *o*, *v*, *x*_0_, *b*, *σ*]. As we will see, each DDM parameter has a specific signature, in terms of its joint impact on these seven quantities. This does not imply however, that different parameter settings necessarily yield distinct moments. In fact, there are changes in the DDM parameters that leave the predicted moments unchanged. This will induce parameter recovery issues, which we will demonstrate later.

**Figure 1:**
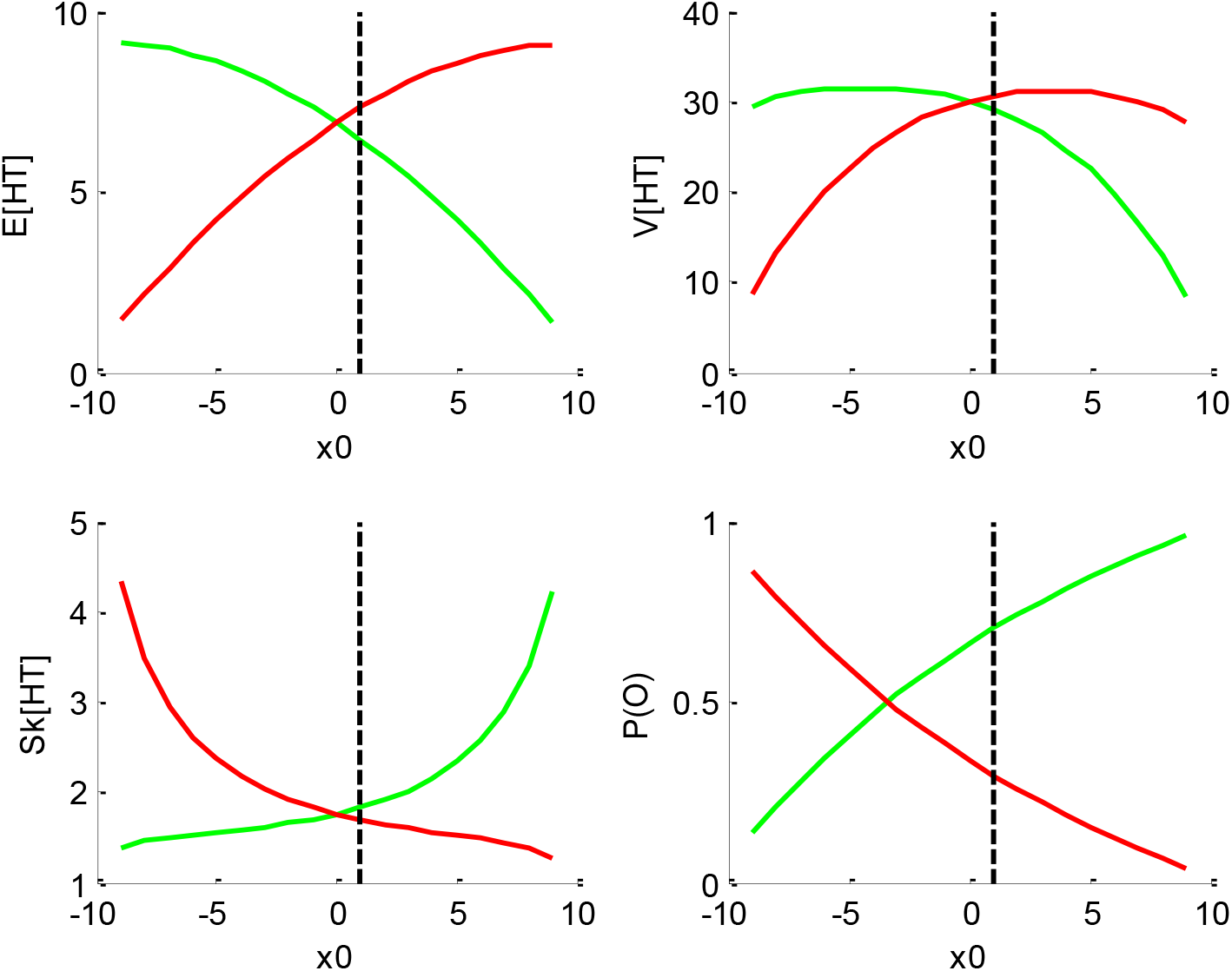
impact of initial bias *x*_0_. In all panels, the colour code indicates the decision outcomes (green: ‘up’ decisions, red: ‘down’ decisions). The black dotted line indicates the default parameter value (for ease of comparison with other figures below). **Upper-left panel**: mean hitting times (y-axis) as a function of initial bias (x-axis). **Upper-right panel**: hitting times’ variance (y-axis) as a function of initial bias (x-axis). **Lower-left panel**: hitting times’ skewness (y-axis) as a function of initial bias (x-axis). **Lower-right panel**: outcome ratios (y-axis) as a function of initial bias (x-axis).

Figure 1 below shows the impact of initial bias *x*_0_.

But let first us summarize the impact of DDM parameters. To do this, we first set model parameters to the following “default” values: *v* = 1/2, *x*_0_ = 1, *b* = 10 and *σ* = 4. This parameter setting yields about 30% decision errors, which we take as a valid reference point for typical studies of decision making. In what follows, we vary each model parameter one by one, keeping the other ones at their default value.

One can see that increasing the initial bias accelerates decision times for ‘up’ decisions, and decelerates decision times for ‘down’ decisions. This is because increasing *x*_0_ mechanically increases the probability of an early upper bound hit, and decreases the probability of an early lower bound hit. Increasing *x*_0_ also decreases (resp., increases) the variance for ‘up’ (resp., ‘down’) decisions, and increases (resp., decreases) the skewness for ‘up’ (resp., ‘down') decisions. Finally, increasing the initial bias increases the ratio of ‘up’ decisions. These are corollary consequences of increasing (resp. decreasing) the probability of an early upper (resp., lower) bound hit. This is because when an increasing proportion of stochastic paths eventually hit a bound very early, this squeezes the distribution of hitting times just above null hitting times. Note that the outcome ratios are not equal to 1/2 when *x*_0_ = 0. This is because the default drift rate *v* is positive, and therefore favours ‘up’ decisions. Most importantly, the initial bias is the only DDM parameter that has opposite effects on HT for ‘up’ and ‘down’ decision outcomes.

Figure 2 below shows the impact of drift rate *v*.

**Figure 2:**
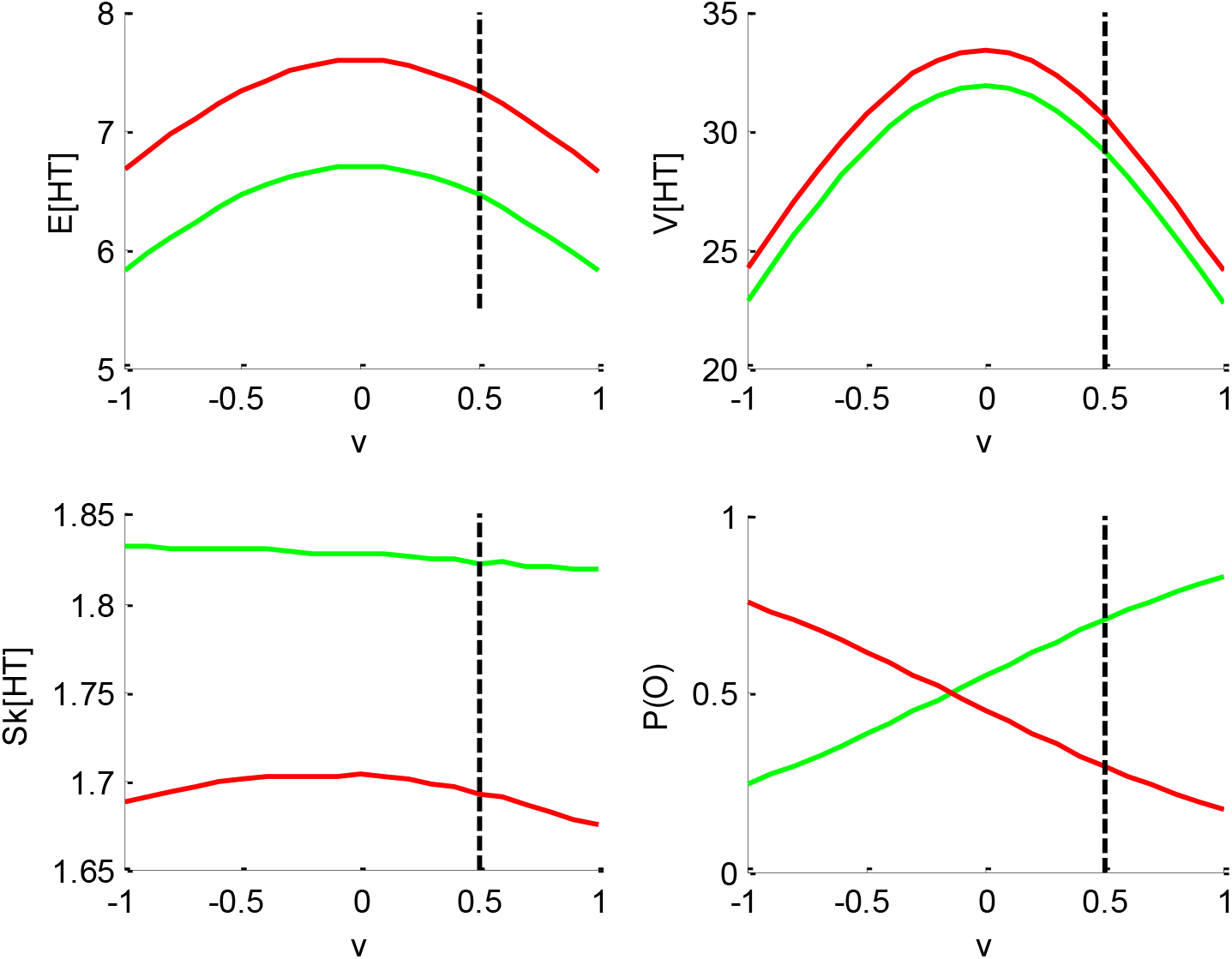
impact of drift rate *v*. Same format as Fig 1.

One can see that the mean and variance of decision times are maximal when the drift rate is null. This is because the probability of an early (upper or lower) bound hit decreases as *v* ⟶ 0. Also, the drift rate has little impact on the HT skewness. Note that, in contrast to the initial bias, the impact of the drift rate on HT is identical for both ‘up’ and ‘down’ decisions. Finally, and as expected, increasing the drift rate increases the ratio of ‘up’ decisions.

Figure 3 below shows the impact of the noise’s standard deviation *σ*.

**Figure 3:**
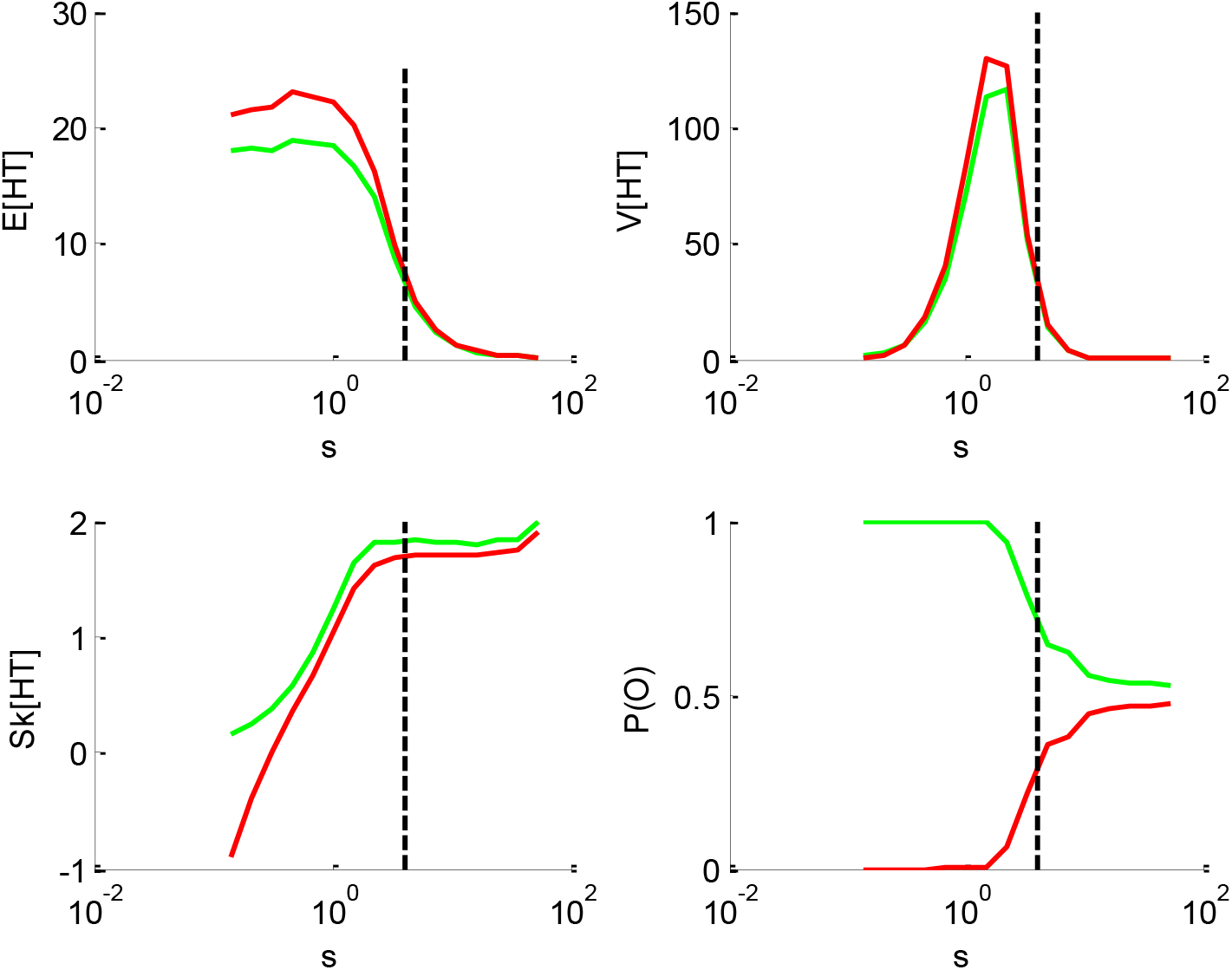
impact of the perturbation’ standard deviation *σ*. Same format as Fig 1 (but the x-axis is now in log-scale).

One can see that increasing the standard deviation decreases the mean HT, and increases its skewness. This is, again, because increasing *σ* increases the probability of an early bound hit. Its impact on the variance, however, is less trivial. When the standard deviation *σ* is very low, increasing *σ* first increases the hitting times’ variance. This is because it progressively frees the system from its deterministic fate, therefore enabling HT variability around the mean. Then, it reaches a critical point above which increasing it further now decreases the variance. This is again a consequence of increasing the probability of an early bound hit. The associated HT squeezing effect can be seen on the skewness, which steadily increases beyond the critical point. Note that the standard deviation has the same impact on HT for ‘up’ and ‘down’ decisions. Finally, increasing the standard deviation eventually maximizes the entropy of the decision outcomes, i.e. *P* (*o*) ⟶ 1/2 when *σ* ⟶ ∞. This is because the relative contribution of the diffusion term eventually masks the drift.

Figure 4 below shows the impact of the bound’s height *b*.

**Figure 4:**
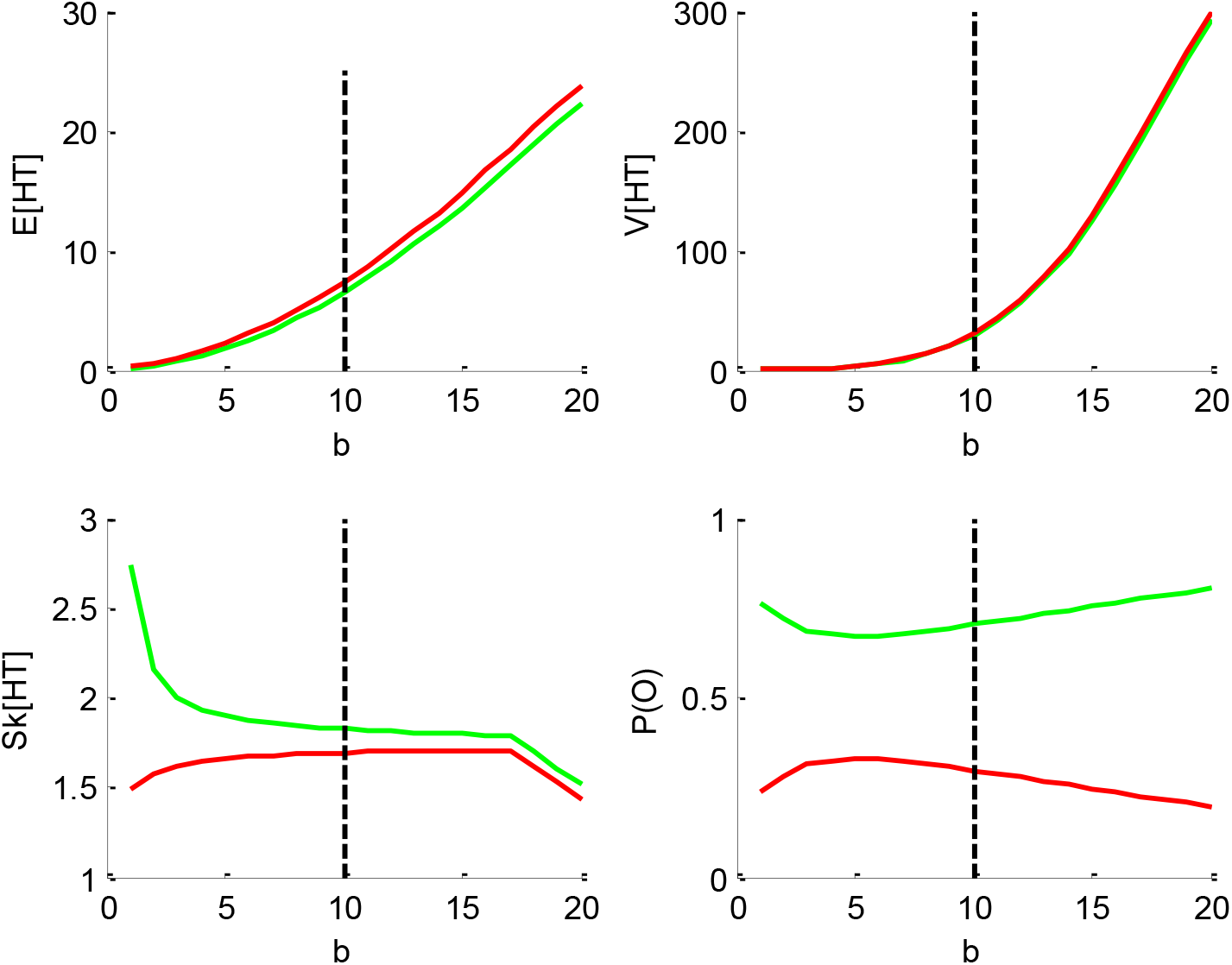
impact of the threshold’s height *b*. Same format as Fig 1.

One can see that increasing the bound’s height increases both the mean and the variance of HT, and decreases its skewness, identically for ‘up’ and ‘down’ decisions. Finally, increasing the threshold’s height decreases the entropy of the decision outcomes, i.e. *P* (*o*) ⟶ 0 or 1 when *b* ⟶ ∞. This directly derives from the fact that increasing *b* decreases the probability of an early bound hit. This effect basically competes with the effect of the standard deviation *σ*, which accelerates HTs. This is why one may say that increasing the threshold’s height effectively increases the demand for evidence strength in favour of one of the decision outcomes.

Note that the impact of the “non-decision” time *T_ND_* simply reduces to shifting the mean of the RT distribution, without any effect on other moments.

In brief, DDM parameters have distinct impacts on the sufficient statistics of response times. This means that they could, in principle, be discriminated from each other. Standard DDM fitting procedures rely on adjusting the DDM parameters so that the RT moments (e.g., up to third order) match model predictions. In what follows, we refer to this as the “method of moments” (see Appendix 2). However, we will see below that the DDM is not perfectly identifiable. One can also see that changing any of these parameters from trial to trial will most likely induce non-trivial variations in RT data. Here, the method of moments may not be optimal, because predictable trial-by-trial variations in DDM parameters will be lumped together with stochastic perturbation-induced variations. One may thus rather attempt to match the trial-by-trial series of raw response times directly with their corresponding first-order moments. In what follows, we refer to this as the “method of trial means” (see Appendix 3). Given the computational cost of deriving expected response times for each trial, this type of approach is typically restricted to the vanilla DDM, since there is no known analytical expression for response time moments under more complex DDM variants.

Below, we suggest a simple and efficient way of performing DDM parameter estimation, which applies to a broad class of DDM variants without significant additional computational burden. This follows from fitting a self-consistency equation that, under a broad class of DDM variants, response times have to obey.

## 3. A self-consistency equation for DDMs

First, note that Equation 2 can be rewritten as follows:

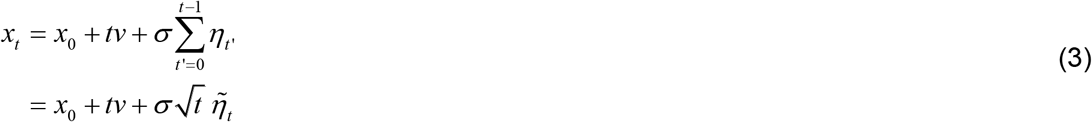

where we coin 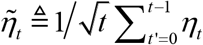 the “normalized cumulative perturbation”. Now let *τ_i_* be the decision time of the *i*^th^ trial. Note that decision times are trivially related to cumulative perturbations because, by definition, 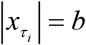. This implies that:

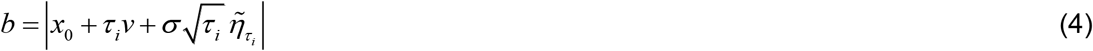

where 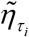 denotes the (unknown) cumulative perturbation term of the *i*^th^ trial.

Information regarding the binary decision outcome *o_i_* ∈ {−1,1} further disambiguates Equation 4 as follows:

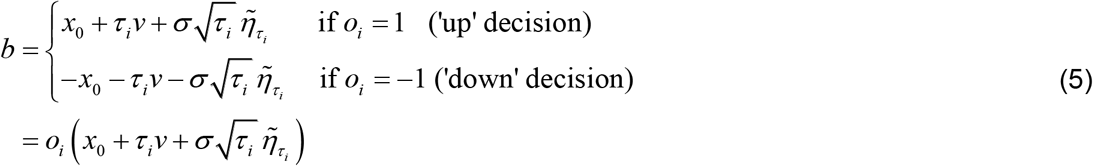

where *o_i_* can only take two possible values (−1 or 1). Equation 5 can then be used to relate decision times directly to DDM model parameters (and cumulative perturbations):

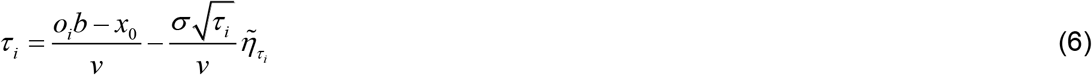

From Equation 6, one can express observed trial-by-trial empirical response times *y_i_* as follows:

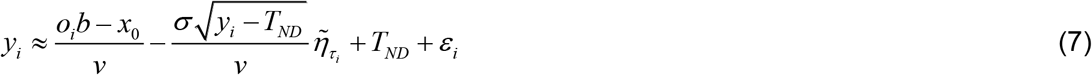

where *ε_i_* are unknown i.i.d. model residuals.

Note that decision times effectively appear on both the left-hand and the right-hand side of Equations 6-7. This is a slightly unorthodox feature, but, as we will see, this has effectively no consequence from the perspective of model inversion. In fact, one can think of Equation 7 as a “self-consistency” constraint that response times have to fulfil under the DDM. This is why we refer to Equation 7 as the *self-consistency equation* of DDMs. This, however, prevents us from using Equation 7 to generate data under the DDM. In other terms, Equation 7 is only useful when analysing empirical RT data.

Figure 5 below exemplifies the accuracy of DDM’s self-consistency equation, using a Monte-Carlo simulation of 200 trials under the vanilla DDM.

**Figure 5:**
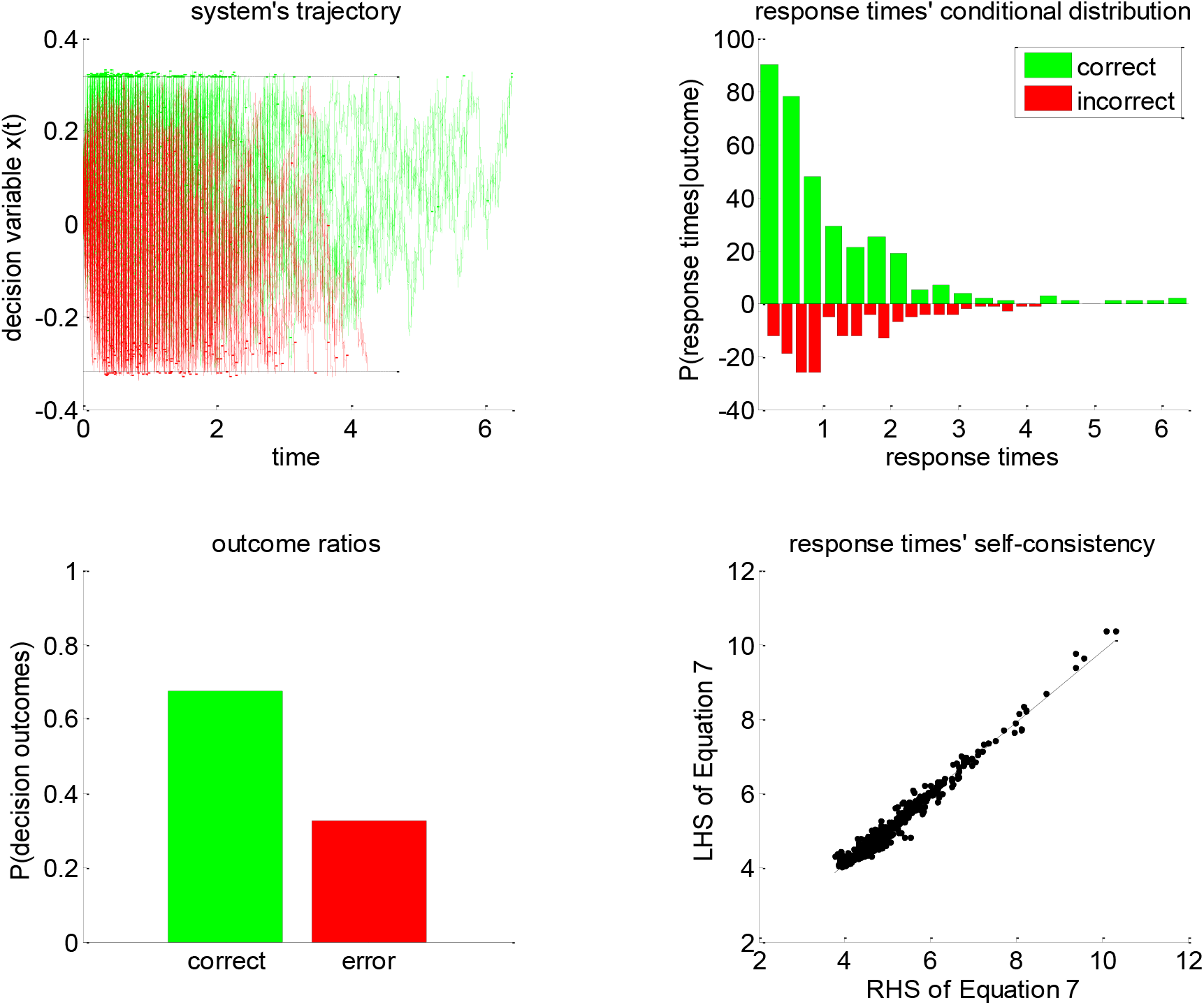
Self-consistency equation. Monte-Carlo simulation of 200 trials of a DDM, with arbitrary parameters (in this example, the drift rate is positive). In all panels, the colour code indicates the decision outcomes, which depends upon the sign of the drift rate (green: correct decisions, red: incorrect decisions). **Upper-left panel**: simulated trajectories of the decision variable (y-axis) as a function of time (x-axis). **Upper-right panel**: response times’ distribution for both correct and incorrect choice outcomes over the 200 Monte-Carlo simulations. **Lower-left panel**: outcome ratios. **Lower-right panel**: the left-hand side of Equation 7 (y-axis) is plotted against the right-hand side of Equation 7 (x-axis), for each of the 200 trials.

One can see that the DDM’s self-consistency equation is valid, i.e. simulated response times almost always equate their theoretical prediction. The few (small) deviations that can be eyeballed on the lower-right panel of Figure 5 actually correspond to simulation artefacts where the decision variable exceeds the bound by some relatively small amount. This happen when the discretization step Δ*t* (cf. Equation 2) is too large when compared to the relative magnitude of the stochastic component of the system’s dynamics. In effect, these artefactual errors grow when *σ*/*v* increases. Nevertheless, in principle, these and other errors would be absorbed in the model residuals *ε*_i_ of Equation 7.

Now recall that recent extensions of vanilla DDMs include e.g., collapsing bounds (Hawkins et al., 2015; Voskuilen et al., 2016) and/or nonlinear transformations of the state-space (Tajima et al., 2016). As the astute reader may have already guessed, the self-consistency equation can be generalized to such DDM variants. Let us assume that Equations 2-3 still hold, i.e. the decision process is still somehow based upon a gaussian random walk. However, we now assume that the decision is triggered when an arbitrary transformation *z*: *x* ⟶ *z*(*x*) of the base random walk *x_t_* has reached a predefined threshold 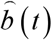 that can vary over time (e.g., a collapsing bound). Equation 5 now becomes:

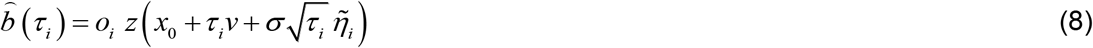

If the transformation *z*: *x* ⟶ *z*(*x*) is invertible (i.e. if *z*^−1^ exists and is unique), then the self-consistency equation for reaction times *y_i_* now generalizes as follows:

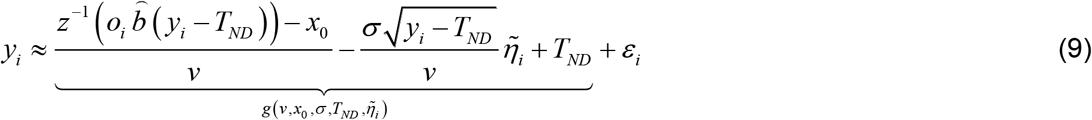

where 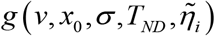 is the “expected” (or rather, “self-consistent”) response time at trial *i*, which depends nonlinearly on DDM parameters (and on response times). Note that one recovers the self-consistency equation of “vanilla” DDM (Equation 7) when setting *z*(*x*) = *z*^−1^ (*x*) = *x* and 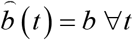.

Importantly, inverting Equation 9 can be used to estimate parameters *γ* and *ω* that control the transformation 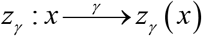 or the collapsing bounds 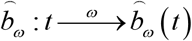, respectively. We will see examples of this in the Results section below. This implies that the self-consistency equation can be used, in conjunction with adequate statistical parameter estimation approaches (see below), for estimating DDM parameters under many different variants of DDM, including those for which no analytical result exists for the response time distribution.

## 4. An overcomplete likelihood approach to DDM inversion

Fitting Equation 9 to response time data reduces to finding the set of parameters that renders the DDM self-consistent. In doing so, normalized cumulative perturbation terms 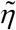 are treated as nuisance model parameters, but model parameters nonetheless. This means that there are more model parameters than there are data points. In other words, Equation 9 induces an “overcomplete” likelihood function 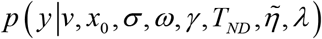

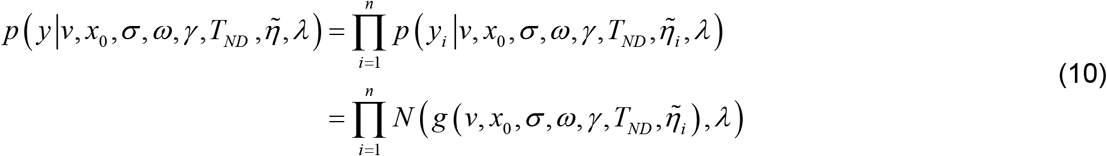

where *λ* is the variance of the model residuals *ε*_*i*_ of Equation 9, *g* (·) is the “self-consistent” response time given in Equation 9, and we have used the (convenient but slightly abusive) notation 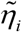 to reference cumulative perturbations w.r.t. to their corresponding trial index.

Dealing with such overcomplete likelihood function requires additional constraints on model parameters: this is easily done within a Bayesian framework. Therefore, we rely on the variational Laplace approach (Daunizeau, 2017; Friston et al., 2007), which was developed to perform approximate bayesian inference on nonlinear generative models (see Appendix 1 for mathematical details). In what follows, we propose a simple set of prior constraints that help regularizing the inference.

### a. Prior moments of the cumulative perturbations: the “no barrier” approximation

Recall that, under the DDM framework, errors can only be due to the stochastic perturbation noise. More precisely, errors are due to those perturbations that are strong enough to deviate the system’s trajectory and make it hit the “wrong” bound (e.g., the lower bound if the drift rate is positive). Let *Q*_=_ be the proportion of correct responses. For example, if the drift rate is positive, then *Q*_=_ corresponds to responses that hit the upper bound. Now let 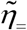 be the critical value of 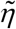 such that 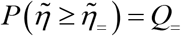 (see Figure 6 below). Then, we know that errors correspond to those perturbations 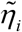 that are smaller than 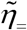. But what do we know about the distribution of perturbations? Importantly, if the DDM’s stochastic evidence accumulation process had no decision bound, then the distribution of normalized cumulative perturbations would be invariant over time and such that 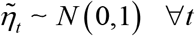. This, in fact, is the very reason why we introduced normalized cumulative perturbations in Equation 3. Under this “no barrier” approximation, one can now derive the conditional expectations 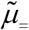 and 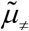 of the perturbation 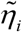, given that the decision outcome *o_i_* is correct or erroneous, respectively:

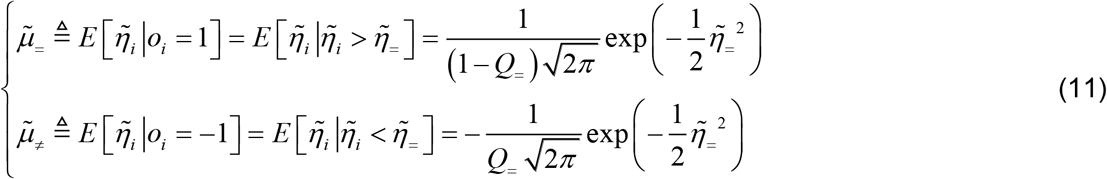

**Figure 6:**
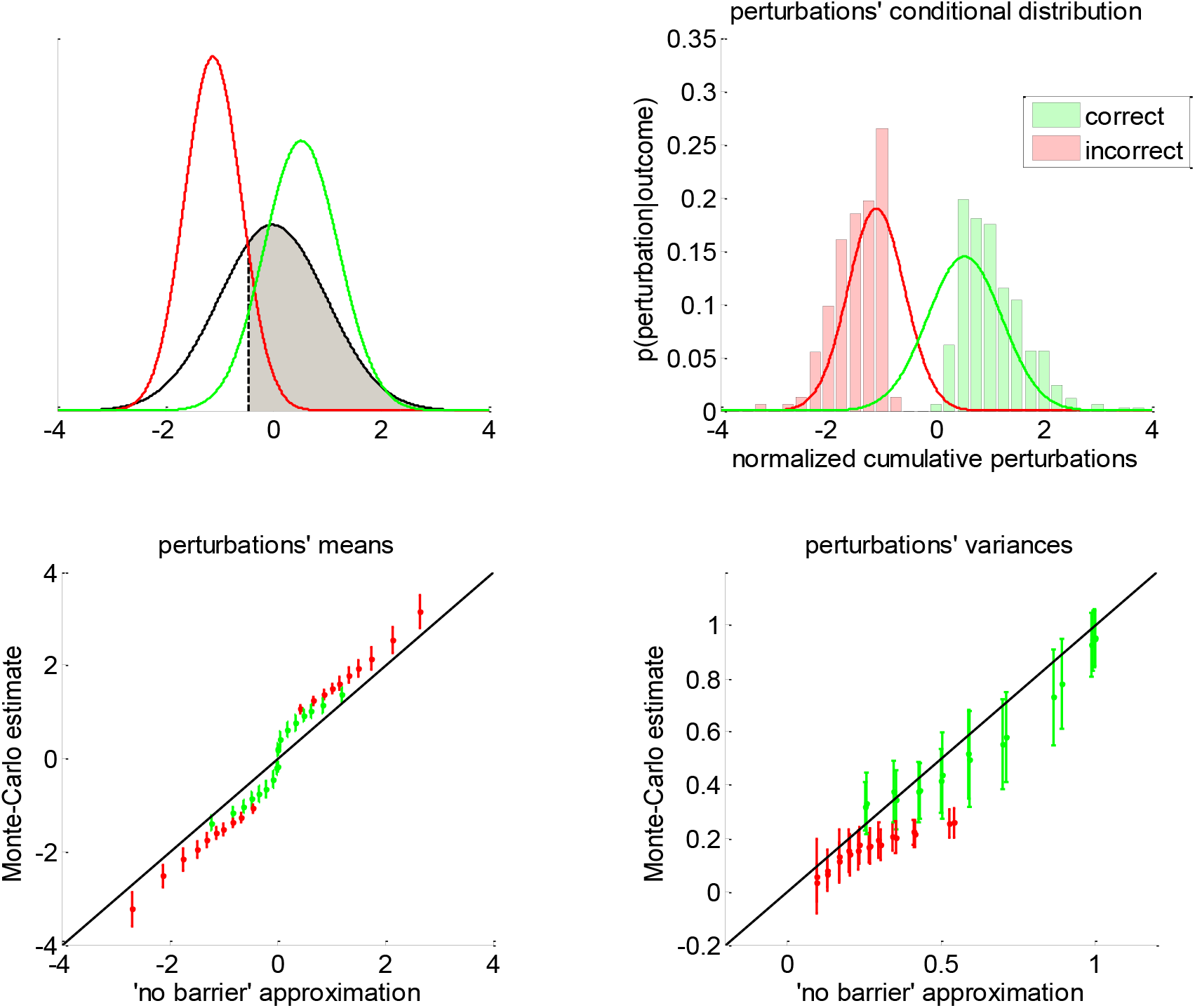
Approximate conditional distributions of the normalized cumulative perturbations. **Upper-left panel**: The black line shows the “no barrier” standard normal distribution of normalized cumulative perturbations. The shaded grey area has size *Q*_=_, and its left bound (dashed black line) is the critical value 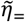 below which cumulative perturbations eventually induce errors. The green and red lines depict the ensuing approximate conditional distributions given in Equation 13. **Upper-right panel**: a representative Monte-Carlo simulation. The green and red bars show the sample histogram of normalized cumulative perturbations for correct and erroneous decisions, respectively (over 200 trials, same simulation as in Figure 5). The green and red lines depict the corresponding approximate conditional normal distributions (cf. Equation 13). **Lower-left panel**: The sample mean estimates of conditional perturbations (y-axis) are plotted against their “no barrier” approximation (x-axis, Equation 11). Monte-Carlo simulations are split according to the sign of the drift rate, and then binned according to deciles of approximate conditional means of normalized cumulative perturbations (green: correct, red: error, errorbars: within-decile means ± standard deviations). The black dotted line shows the identity mapping (perfect approximation). **Lower-right panel**: The sample variance estimates of normalized cumulative perturbations (y-axis) are plotted against their “no barrier” approximation (x-axis, Equation 12). Same format as lower-left panel.

Equation 11 is obtained from the known expression of first-order moments of a truncated normal density *N* (0,1). Critically, Equation 11 does not depend upon DDM parameters. Of course, the same logic extends to conditional variances 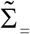 and 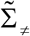, whose analytical expressions are given by:

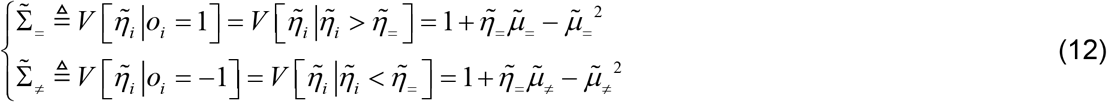

A simple moment-matching approach thus suggests to approximate the conditional distribution 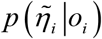 of normalized cumulative perturbations as follows:

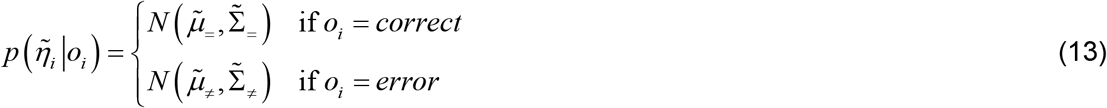

where the correct/error label depends on the sign of the drift rate. This concludes the derivation of our simple “no barrier” approximation to the conditional moments of cumulative perturbations.

Note that we derived this approximation without accounting for the (only) mathematical subtlety of the DDM: namely, the fact that decision bounds formally act as ‘absorbing barriers’ for the system (Broderick et al., 2009). Critically, absorbing barriers induce some non-trivial forms of dynamical degeneracy. In particular, they eventually favour paths that are made of extreme samples of the perturbation noise. This is because these have a higher chance of crossing the boundary, despite being comparatively less likely than near-zero samples under the corresponding “no barrier” distribution. One may thus wonder whether ignoring absorbing barriers may invalidate the moment-matching approximation given in Equations 11-13. To address this concern, we conducted a series of 1000 Monte-Carlo simulations, where DDM parameters were randomly drawn (each simulation consisted of 200 trials of the same decision system). We use these to compare the sample estimates of first-and second-order moments of normalized cumulative perturbations and their analytical approximations (as given in Equations 11-12). The results are given in Figure 6 below.

One can see on the upper-right panel of Figure 6 that the distribution of normalized cumulative perturbations may strongly deviate from the standard normal density. In particular, this distribution clearly exhibits two modes, which correspond to correct and incorrect decisions, respectively. We have observed this bimodal shape across almost all Monte-Carlo simulations. This means that bound hits are less likely to be caused by perturbations of small magnitude than expected under the “no-barrier” distribution (cf. lack of probability mass around zero). Nevertheless, the ensuing approximate conditional distributions seem to be reasonably matched with their sample estimates. In fact, lower panels of Figure 6 demonstrate that sample means and variances of normalized cumulative perturbations are well approximated by Equations 11-12 for a broad range of DDM parameters. We note that the “no-barrier” approximation tends to slightly underestimate first-order moments, and overestimate second-order moments. This bias is negligible however, when compared to the overall range of variations of conditional moments. In brief, the effect of absorbing barriers on system dynamics has little impact on the conditional moments of normalized cumulative perturbations.

When fitting the DDM to empirical RT data, one thus wants to enforce the distributional constraint in Equations 11-13 onto the perturbation term in Equation 9. This can be done using a change of variable 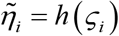, where *ς* are non-constrained dummy variables and *h*: *ς_i_* ⟶ *h* (*ς_i_*) is the following moment-enforcing mapping:

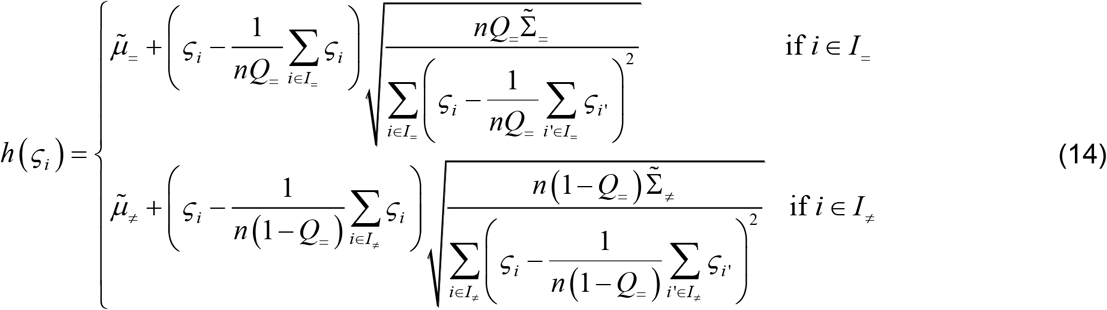

where *I*_=_ and *I*_≠_ are the indices of correct and incorrect trials, respectively (and *n* is the total number of trials). Equation 14 ensures that the sample moments of the estimated normalized cumulative perturbations 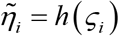 match Equations 11-12, irrespective of the dummy variable *ς*. This also implies that the effective degrees of freedom of the constrained model are in fact lower than what the native self-efficacy function would suggest.

### b. Prior constraints on native DDM parameters

In addition, one may want to introduce the following prior constraints on the native DDM parameters:

- The bound’s height *b* is necessarily positive. This positivity constraint can be enforced by replacing *b* with a non-bounded parameter *φ*_1_, which relates to the bound’s height through the following mapping: *b* = exp(*φ*_1_). We note that parameters *ω*of collapsing bounds 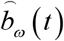 may not have to obey such positivity constraint.
- The standard deviation *σ* is necessarily positive. Again, this can be enforced by replacing it with the following mapped parameter *φ*_2_: *σ* = exp(*φ*_2_).
- The non-decision time *T_ND_* is necessarily positive and smaller than the minimum observed reaction time. This can be enforced by replacing the native non-decision time with the following mapped parameter *φ*_3_: *T*_*ND*_ = min (*RT*)*s* (*φ*_3_), where *s* (•) is the standard sigmoid mapping.
- The initial bias *x*_0_ is necessarily constrained between −*b* and *b*. This can be enforced by replacing the native initial condition with the following mapped parameter *φ*_4_: *x*_0_ = exp(*φ*_1_)(2*s* (*φ*_4_) −1).
- In principle, the drift rate *v* can be either positive or negative. However, its magnitude is necessarily smaller than 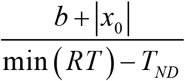, which corresponds to its “ballistic” limit (see Appendix 6 for more details). This can be enforced by replacing the native drift rate with the following mapped parameter φ_5_: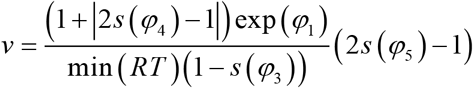

Here again, we use the set of dummy variables *φ*_1:5_ in lieu of native DDM parameters. The statistical efficiency of the ensuing overcomplete approach can be evaluated by simulating RT and choice data under different settings of the DM parameters, and then comparing estimated and simulated parameters. Below, we use such recovery analysis to compare the overcomplete approach with standard DDM fitting procedures.

### c. Accounting for predictable trial-by-trial RT variability

Critically, the above overcomplete approach can be extended to ask whether trial-by-trial variations in DDM parameters explain trial-by-trial variations in observed RT, above and beyond the impact of the random perturbation term in Equation 7. For example, one may want to assess whether predictable variations in e.g., the drift term, accurately predict variations in RT data. This kind of questions underlies many recent empirical studies of human and/or animal decision making. In the context of perceptual decision making, the drift rate is assumed to derive from the strength of momentary evidence, which is controlled experimentally and varies in a trial-by-trial fashion (Bitzer et al., 2014; Huk and Shadlen, 2005). A straightforward extension of this logic to value-based decisions implies that the drift rate should vary in proportion to the value difference between alternative options (De Martino et al., 2012; Krajbich et al., 2010; Lopez-Persem et al., 2016). In both cases, a prediction for drift rate variations across trials is available, which is likely to induce trial-by-trial variations in choice and RT data. Let *D* be a known predictor variable, which is expected to capture trial-by-trial variations in some DDM parameter (e.g., the drift rate). One may then alter the self-consistency equation such that DDM parameters are treated as affine functions of trial-by-trial predictors (e.g., 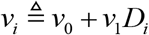), and exploit trial-by-trial variations in response times to fit the ensuing offset and slope parameters (here, *v*_0_ and *v*_1_). Alternatively, one can simply set the drift rate to the predictor variable (i.e. assume *a priori v*_0_ = 0 and *v*_1_ = 1), which is currently the favourite approach in the field. As we will see below, this significantly improves model identifiability for the remaining parameters. This is because trial-by-trial variations in the drift rate will only accurately predict trial-by-trial variations in response time data if the remaining parameters are correctly set. This is just an example of course, and one can see how easily any prior dependency to a predictor variable could be accounted for. The critical point here is that the overcomplete approach can exploit predictable trial-by-trial variations in RT data to improve the inference on model parameters.

## 5. Parameter recovery analysis

In what follows, we use numerical simulations to evaluate the approach’s ability to recover DDM parameters. Our parameter recovery analyses proceed as follows. First, we sample 1000 sets of model parameters *φ*_1:5_ under some arbitrary distribution. Second, for each of these parameter, we simulate a series of N=200 DDM trials according to Equation 2 above. Third, we fit the DDM to each series of simulated reaction times (200 data points) and extract parameter estimates. Last, we compare simulated and estimated parameters to each other. In particular, we measure the relative estimation error for each DDM parameter. We also quantify potential non-identifiability issues using so-called recovery matrices and the ensuing identifiability index. We note that details regarding parameter recovery analyses can be found in the Appendix 4 of this manuscript (along with definitions of the relative estimation error *REE*, recovery matrices and identifiability index Δ*V*).

To begin with, we will focus on “vanilla” DDMs, because they provide a fair benchmark for parameter estimation methods. In this context, we will compare the overcomplete approach with two established methods (Boehm et al., 2018, 2018; Moens and Zenon, 2017), namely: the “method of moments” and the “method of trial means”. These methods are summarized in Appendix 2 and 3, respectively. In brief, the former attempts to match empirical and theoretical moments of RT data. We expect this method to perform best when DDM parameters are fixed across trials. The latter rather attempts to match raw trial-by-trial RT data to trial-by-trial theoretical RT means. This will be most reliable when DDM parameters (e.g., the drift rate) vary over trials. Note that, in all cases, we inserted the prior constraints on DDM parameters given in the Section 3.b above, along with standard normal priors on unmapped parameters *φ*_1:5_. We will therefore compare the ability of these methods to recover DDM parameters (i) when no parameter is fixed (full parameter set), (ii) when the drift rate is fixed, and (iii) when drift rates vary over trials.

Finally, we perform a parameter recovery analysis in the context of a generalized DDM, which includes collapsing bounds. This will serve to demonstrate the flexibility and robustness of the overcomplete approach.

### a. Vanilla DDM: recovery analysis for the full parameter set

First, we compare the three approaches when all DDM parameters have to be estimated. This essentially serves as a reference point for the other recovery analyses. The ensuing recovery analysis is summarized in Figure 7 below, in terms of the comparison between simulated and estimated parameters.

**Figure 7:**
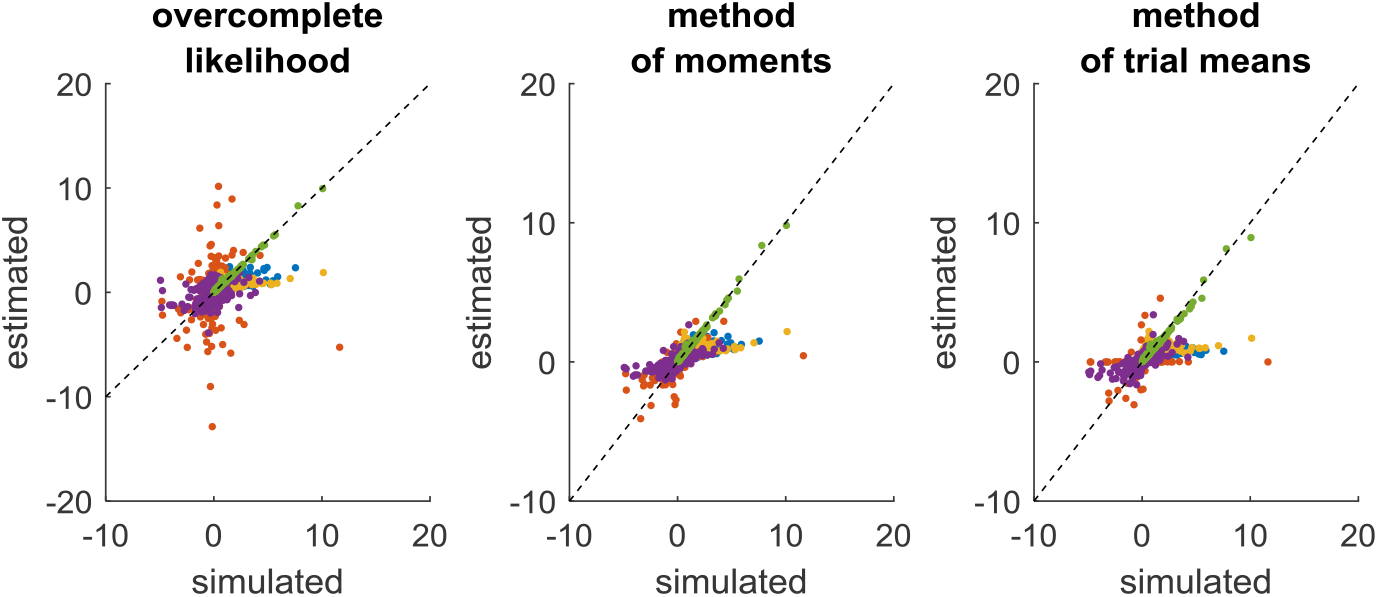
Comparison of simulated and estimated DDM parameters (full parameter set). **Left panel**: estimated parameters using the overcomplete approach (y-axis) are plotted against simulated parameters (x-axis). Each dot is a Monte-Carlo simulation and different colors indicate distinct parameters (blue: *σ*, red: *v*, yellow: *b*, purple: *x*_0_, green: *T_ND_*). The black dotted line indicate the identity line (perfect estimation). **Middle panel**: method of moments, same format as left panel. **Right panel**: method of trial means, same format as left panel.

Unsurprisingly, parameter estimates depend on the chosen estimation method, i.e. different methods exhibit distinct estimation errors structures. In addition, estimated and simulated parameters vary with similar magnitudes, and no systematic estimation bias is noticeable. It turns out that, in this setting, estimation error is minimal for the method of moments, which exhibits lower error than both the overcomplete approach (mean error difference: Δ log(*REE*) = 0.27 ± 0.03, p<10^−4^, two-sided F-test) and the method of moments (mean error difference: Δ log(*REE*) = 0.26 ± 0.02, p<10^−4^, two-sided F-test). However, the overcomplete approach and the method of trial means yield comparable estimation errors (mean error difference: Δ log (*REE*) = 0.006 ± 0.04, p=0.88, two-sided F-test).

Now, although estimation errors enable a coarse comparison of methods, it does not provide any quantitative insight regarding potential non-identifiability issues. We address this using recovery matrices (see Appendix 4), which are shown on Figure 8 below.

**Figure 8:**
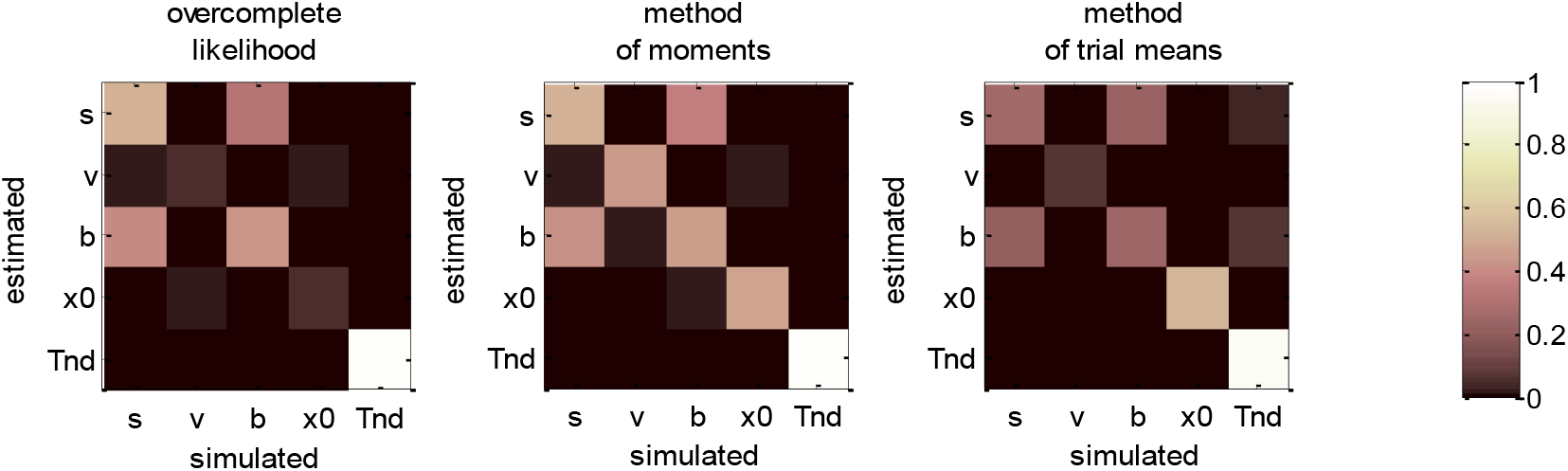
DDM parameter recovery matrices (full parameter set). **Left panel**: overcomplete approach. **Middle panel**: method of moments. **Right panel**: Method of trial means. Each line shows the squared partial correlation coefficient between a given estimated parameter and each simulated parameter (across 1000 Monte-Carlo simulations). Note that perfect recovery would exhibit a diagonal structure, where variations in each estimated parameter is only due to variations in the corresponding simulated parameter. Diagonal elements of the recovery matrix measure “correct estimation variability”, i.e. variations in the estimated parameters that are due to variations in the corresponding simulated parameter. In contrast, non-diagonal elements of the recovery matrix measure “incorrect estimation variability”, i.e. variations in the estimated parameters that are due to variations in other parameters. Strong non-diagonal elements in recovery matrices thus signal pairwise non-identifiability issues.

None of the estimation methods is capable of perfectly identifying DDM parameters (except *T_ND_*), i.e. all methods exhibit strong non-identifiability issues. In particular, variations in the perturbations’ standard deviation *σ* are partially confused with variations in the bound’s height *b*, and reciprocally. This is because increasing both at the same time leaves RT trial-by-trial variability unchanged. Therefore, RT produced under strong neural perturbations can be equally well explained with a small bound height (and reciprocally). Interestingly, drift rate estimates are the least reliable: though their amount of “correct variability” is decent for the method of moments (45.3%), it is very low for both the overcomplete approach (5.3%) and the method of trial means (7.5%). If anything, non-identifiability issues are strongest for the overcomplete approach, which also exhibits weak “correct variability” for initial conditions (5.1%).

### b. Vanilla DDM: recovery analysis with a fixed drift rate

In fact, we expect non-identifiability issues of this sort, which were already highlighted in early DDM studies (Ratcliff, 1978). Note that this basic form of non-identifiability is easily disclosed from the self-consistency equation, which is invariant to a rescaling of all DDM parameters (except *T_ND_*. In other terms, response times are left unchanged if all these parameters are rescaled by the same amount. Although this problematic invariance would disappear if a single DDM parameter was fixed rather than fitted, other non-identifiability issues may still hamper DDM parameter recovery. To test this, we re-performed the above parameter recovery analysis, but this time informing estimation methods about the drift rate, which was set to its simulated value. We note that such arbitrary reduction of the parameter space is routinely performed, as it was already suggested in seminal empirical applications of the DDM (Ratcliff, 1978). Figure 9 below summarizes the ensuing comparison between simulated and estimated parameters.

**Figure 9:**
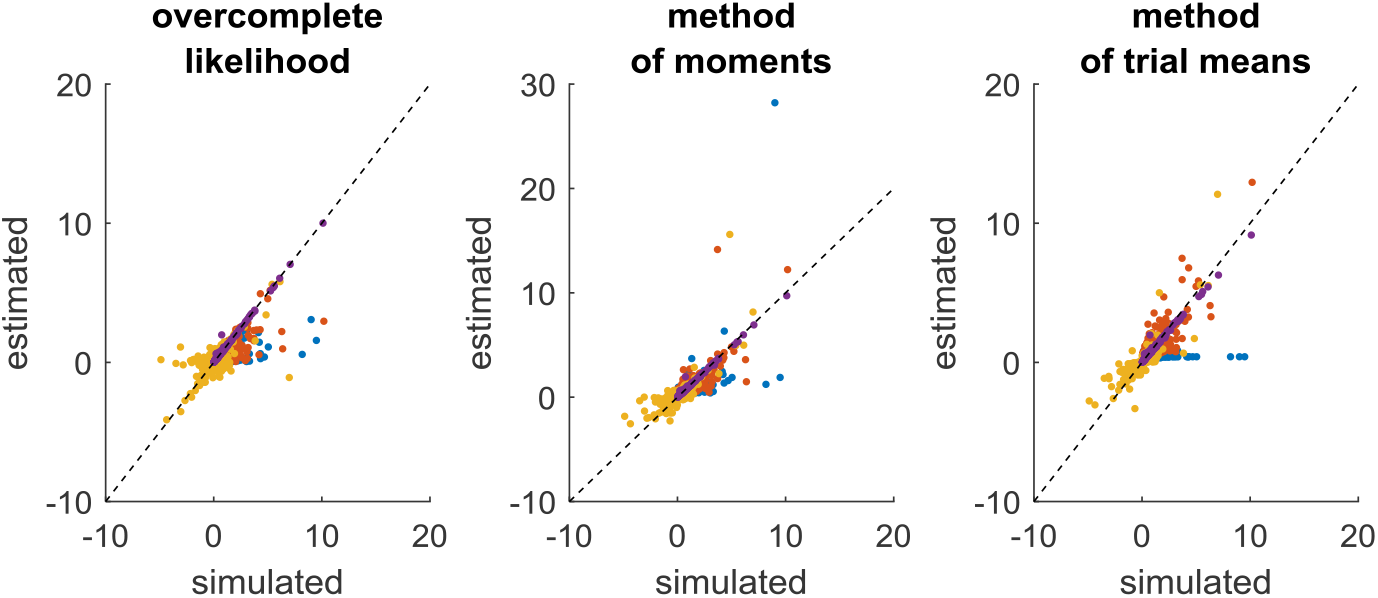
Comparison of simulated and estimated DDM parameters (fixed drift rates). Same format as Fig.7, except for the color code in upper panels (blue: *σ*, red: *b*, yellow: *x*_0_, purple: *T_ND_*).

Comparing Figure 7 and 9 provides a clear insight regarding the impact of reducing the DDM’s parameter space. In brief, estimation errors decrease for all methods, which seem to provide much more reliable parameter estimates. The method of moments still yields the most reliable parameter estimates, eventually exhibiting lower error than the overcomplete approach (mean error difference: Δ log (*REE*) = 0.21 ± 0.03, p=0.04, two-sided F-test) and the method of trial means (mean error difference: Δ log(*REE*) = 0.53 ± 0.03, p<10^−4^, two-sided F-test). In addition, the overcomplete approach yields lower estimation error than the method of trial means (mean error difference: Δ log(*REE*) = 0.33 ± 0.04, p<10^−4^, two-sided F-test). The reason why the methods of trial means performs worst here is that it is blind to trial-by-trial variability in the data (beyond mean RT differences between the two decision outcomes). This is not the case however, for the two other methods.

We then evaluated non-identifiability issues using recovery matrices, which are summarized in Figure 10 below.

**Figure 10:**
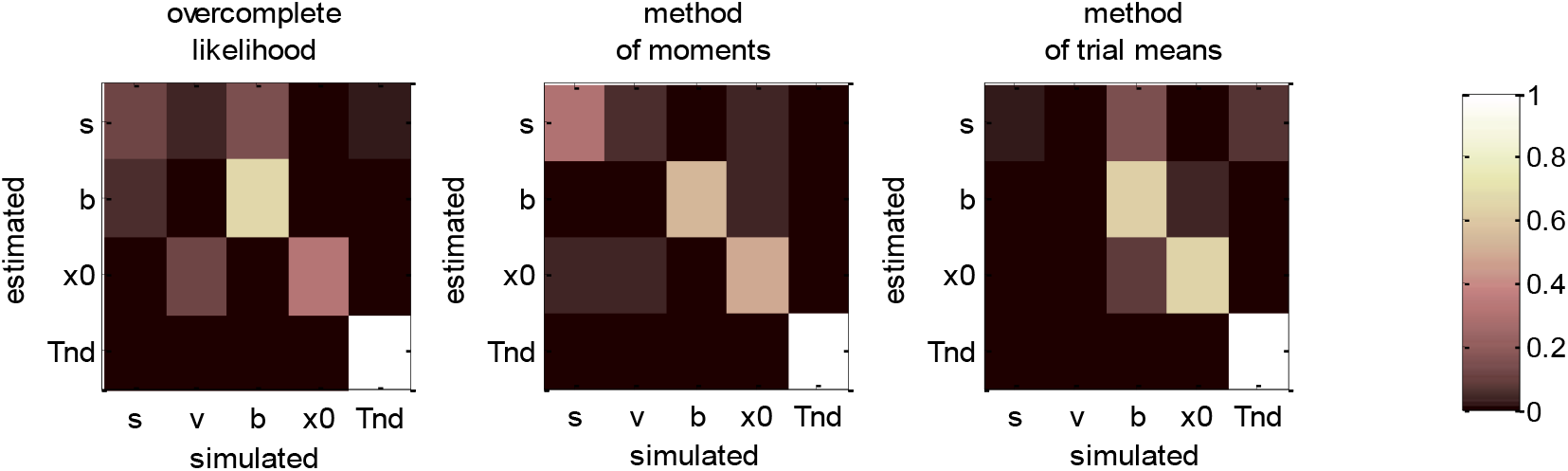
DDM parameter recovery matrices (fixed drift rates). Same format as Fig. 8, except that recovery matrices do not include the line that corresponds to the drift rate estimates. Note, however, that we still account for variations in the remaining estimated parameters that are attributable to variations in simulated drift rates.

Figure 10 clearly demonstrates an overall improvement in parameter identifiability (compare to Figure 8). In brief, most parameters are now identifiable, at least for the method of moments (which clearly performs best) and the overcomplete approach. Nevertheless, some weaker non-identifiability issues still remain, even when fixing the drift rate to its simulated value. For example, the overcomplete approach and the method of trial means still somehow confuse bound’s heights with perturbations’ standard deviations. More precisely, 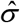 shows unacceptably weak “correct variations” (overcomplete approach: 12.3%, method of trial means: 2.7%), when compared to “incorrect variations” due to the bound’s height (overcomplete approach: 12.4%, method of trial means: 14.3%). Note that this does not hold for the method of moments, for which 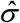 shows strong “correct variations” (30.2%). Having said this, even the method of moments exhibit partial non-identifiability issues, in particular between perturbations’ standard deviations and drift rates (incorrect variations: 4.1%).

We note that fixing another DDM parameter, e.g. the noise’s standard deviation *σ*(instead of *v*), would not change the relative merits of estimation methods in terms of parameter recovery. In other words, the above results are representative of the impact of fixing any DDM parameter. But situations where the drift rate is fixed can be directly compared with situations where one is attempting to exploit predictable drift rates trial-by-trial variations, which is the focus of the next section.

### c. Vanilla DDM: recovery analysis with varying drift rates

Now, accounting for predictable trial-by-trial variations in model parameters may, in principle, improve model identifiability. This is due to the fact that the net effect of each DDM parameter depends upon the setting of other parameters. Let us assume, for example, that the drift rate varies across trials according to some predictor variable (e.g., the relative evidence strength of alternative options in the context of perceptual decision making). The impact of other DDM parameters will not be the same, depending on whether the drift rate is high or low. In turn, there are fewer settings of these parameters that can predict trial-by-trial variations in RT data from variations in drift rate. To test this, we re-performed the recovery analysis, this time setting the drift rate according to a varying predictor variable, which is supposed to be known. The ensuing comparison between simulated and estimated parameters is summarized in Figure 11 below.

**Figure 11:**
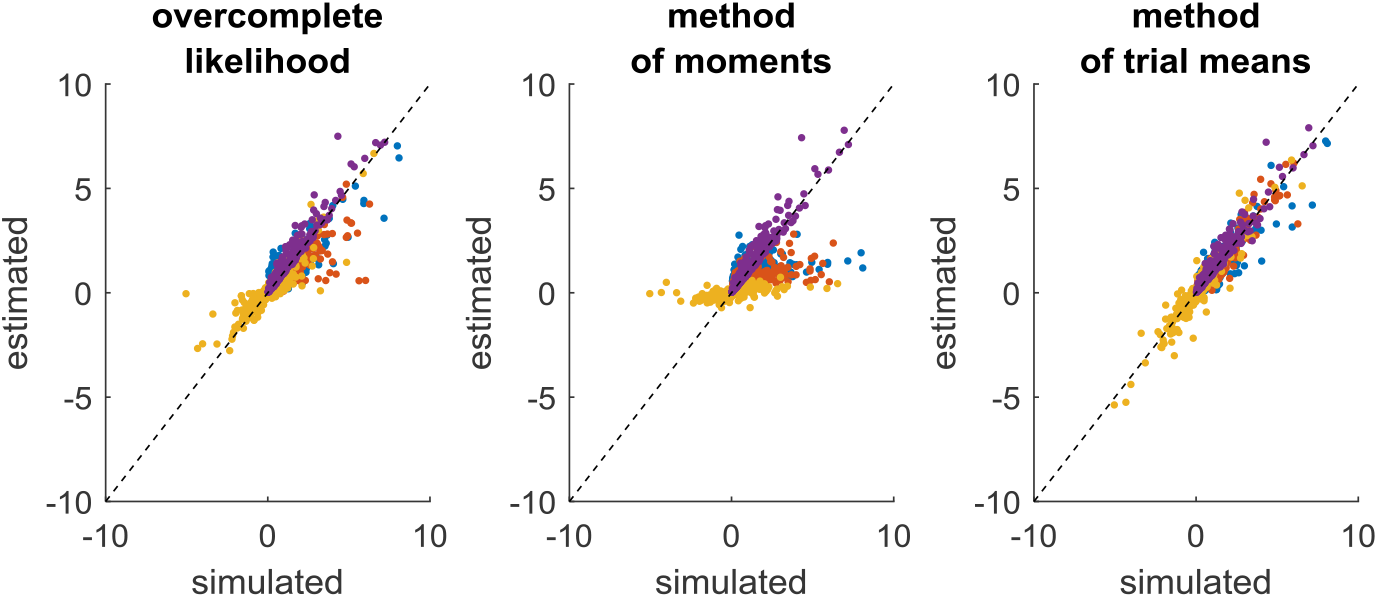
Comparison of simulated and estimated DDM parameters (varying drift rates). Same format as Fig.9.

On the one hand, the estimation error has now been strongly reduced, at least for the overcomplete approach and the method of trial means. On the other hand, estimation error has increased for the method of moments. This is because the method of moments confuses trial-by-trial variations that are caused by variations in drift rates with those that arise from the DDM’s stochastic “neural” perturbation term. This is not the case for the overcomplete approach and the method of trial means. In turn, the method of moments now shows much higher estimation error than the overcomplete approach (mean error difference: Δ log(*REE*) = 0.55 ± 0.03, p<10^−4^, two-sided F-test) or the method of trial means (mean error difference: Δ log(*REE*) = 0.83 ± 0.04, p<10^−4^, two-sided F-test). Note that the latter eventually performs slightly better than the overcomplete approach (mean error difference: Δ log(*REE*) = 0.28 ± 0.03, p=0.04, two-sided F-test).

Figure 12 below then summarizes the evaluation of non-identifiability issues, in terms of recovery matrices.

**Figure 12:**
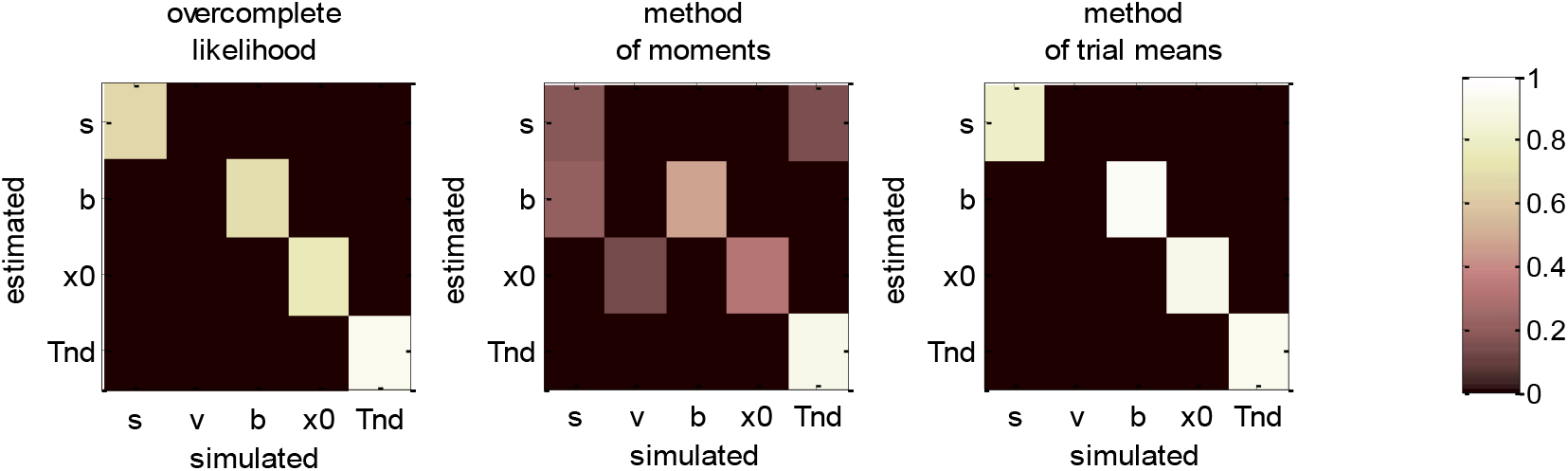
DDM parameter recovery matrices (varying drift rates). Same format as Fig. 10, except that fixed drift rates are replaced by their average across DDM trials.

For the overcomplete approach and the method of trial means, Figure 12 shows a further improvement in parameter identifiability (compare to Figures 8 and 10). For these two methods, all parameters are now well identifiable (“correct variations” are always greater than 67.2% for all parameters), and no parameter estimate is strongly influenced by other simulated parameters. This is a simple example of the gain in statistical efficiency that result from exploiting known trial-by-trial variations in DDM model parameters. The situation is quite different for the method of moments, which exhibits clear non-identifiability issues for all parameters except the non-decision time. In particular, the bound’s height is frequently confused with the perturbations’ standard deviation (20.3% of “incorrect variations”), the estimate of which has become unreliable (only 17.6% of “correct variations”).

We note that the gain in parameter recovery that obtains from exploiting predictable trial-by-trial variations in drift rates (with either the method of trial means or the overcomplete approach) does not generalize to situations where drift rates are defined in term of an affine transformation of some predictor variable (see section 4.c above). This is because the ensuing offset and slope parameters would then need to be estimated along with other native DDM parameters. In turn, this would reintroduce identifiability issues similar or worse than when the full set of parameters have to be estimated (cf. section 4.a). This is why people then typically fix another DDM parameter, e.g. the standard deviation *σ*(Ratcliff et al., 2016). But the risk of drawing erroneous conclusions, e.g. blindly interpreting differences due to *σ* in terms of differences in other DDM parameters, should invite modelers to be cautious with this kind of strategy.

### d. Generalized DDM: recovery analysis with collapsing bounds

We now consider generalized DDMs that include collapsing bounds. More precisely, we will consider a DDM where the bound 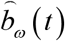 is exponentially decaying in time, i.e.: 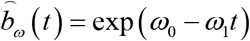, where *ω*_0_ and *ω*_1_ control the bound’s initial height and decay rate, respectively. This DDM variant reduces to the vanilla DDM when *ω*_1_ ≈ 0, in which case the parameter *ω*_0_ is formally identical to the vanilla bound’s height *b*. When *ω*_1_ ≠ 0 however, collapsing bounds induce a causal dependency between choice accuracy and response times that cannot be captured by the vanilla DDM (Hawkins et al., 2015; Tajima et al., 2016; Voskuilen et al., 2016; Zhang, 2012; Zhang et al., 2014).

In what follows, we report the results of a recovery analysis, in which data was simulated under the above generalized DDM (with drift rates varying across trials). We note that, under such generalized DDM variant, no analytical solution is available to derive RT moments. Applying the method of moments or the method of trial means to such generalized DDM variant thus involves either sampling schemes or numerical solvers for the underlying Fokker-Planck equation (Shinn et al., 2020). However, the computational cost of deriving trial-by-estimates of RT moments precludes routine data analysis using these methods, which is why most model-based studies are currently restricted to the vanilla DDM (Fengler et al., 2020). In turn, we do not consider here such computationally intensive extensions of the method of moments and/or method of trial means. In this setting, they thus do not rely on the correct generative model. The ensuing estimation errors and related potential identifiability issues should thus be interpreted in terms of the (lack of) robustness against simplifying modelling assumptions. This is not the case for the overcomplete approach, which bypasses this computational bottleneck and hence generalizes without computational harm to such DDM variants.

Figure 13 below summarizes the ensuing comparison between simulated and estimated parameters.

**Figure 13:**
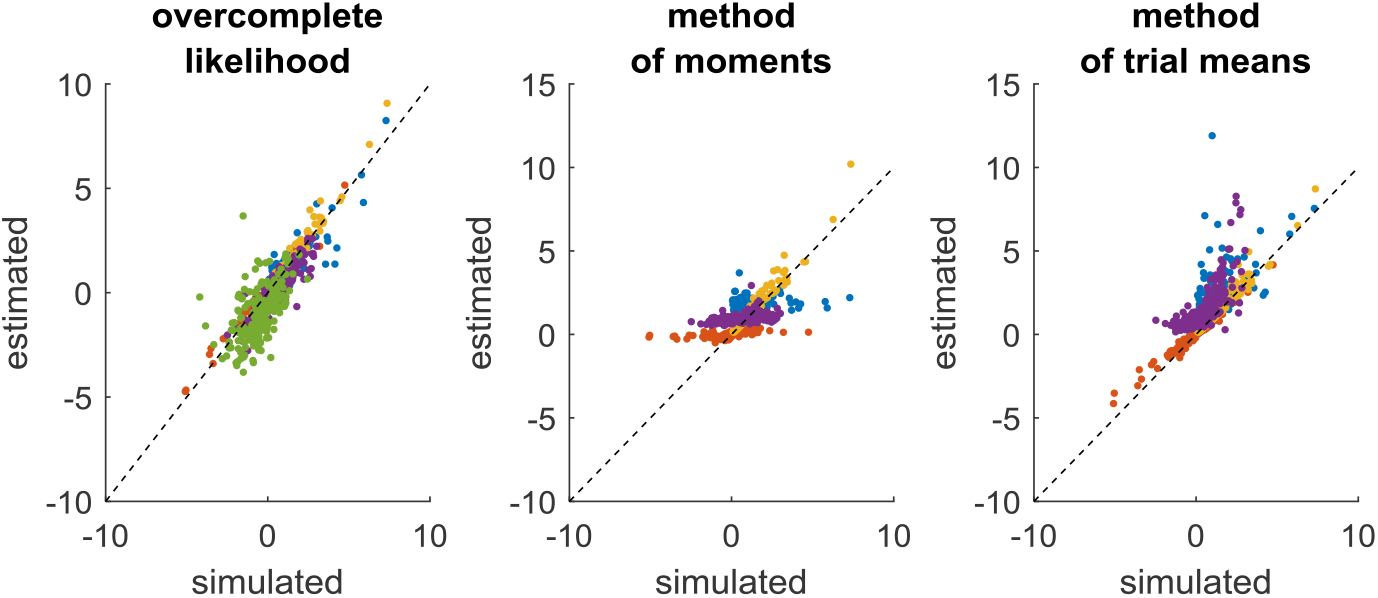
Comparison of simulated and estimated DDM parameters (collapsing bounds). Same format as Fig.9, except that the left panel includes an additional parameter (*w*_1_: green color), which controls the decay rate of DDM bounds.

In brief, the overcomplete approach seems to perform as well as for non-collapsing bounds (see Figure 11). Expectedly however, the method of moments and the method of trial means do incur some reliability loss. Quantitatively, the overcomplete approach shows much smaller estimation error than the method of moments (mean error difference: Δ log(*REE*) = 0.88 ± 0.05, p<10^−4^, two-sided F-test) or the method of trial means (mean error difference: Δ log(*REE*) = 0.61± 0.05, p<10^−4^, two-sided F-test).

Figure 14 below then summarizes the ensuing evaluation of non-identifiability issues, in terms of recovery matrices.

**Figure 14:**
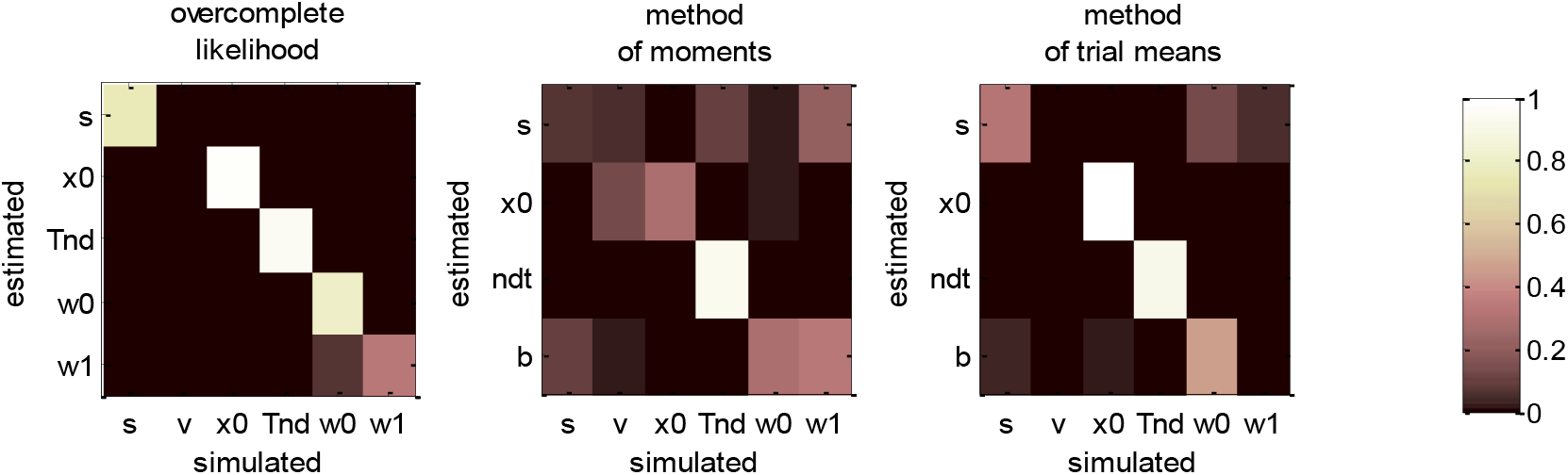
DDM parameter recovery matrices (collapsing bounds). Same format as Fig. 12, except that recovery matrices now also include the bound’s decay rate parameter (*w*_1_), in addition to the bound’s initial height (*w*_0_).

For the overcomplete approach, Figure 14 shows a similar parameter identifiability than Figure 12. In brief, all parameters of the generalized DDM are identifiable from each other (the amount of “correct variations” is 33.8% for the bound’s decay parameter, and greater than 75.5% for all other parameters). This implies that including collapsing bounds does not impact parameter recovery with this method. This is not the case for the two other methods, however. In particular, the method of moments confuses the perturbations’ standard deviation with the bound’s decay rate (7.2% “correct variations” against 20.8% “incorrect variations”). This is also true, though to a lesser extent, for the method of trial means (31.6% “correct variations” against 5.4% “incorrect variations”). Again, these identifiability issues are expected, given that neither the method of moments nor the method of trial means (or, more properly, the variant that we use here) rely on the correct generative model. Maybe more surprising is the fact that these methods now exhibit non-identifiability issues w.r.t. parameters that they can, in principle, estimate. This exemplifies the sorts of interpretation issues that arise from relying on methods that neglect decision-relevant mechanisms. We will comment on this and related issues further in the Discussion section below.

### e. Summary of recovery analyses

Figure 15 below summarizes all our recovery analyses above, in terms of the average (log-) relative estimation error *REE* and the parameter identifiability index Δ*V* (cf. Appendix 4).

**Figure 15:**
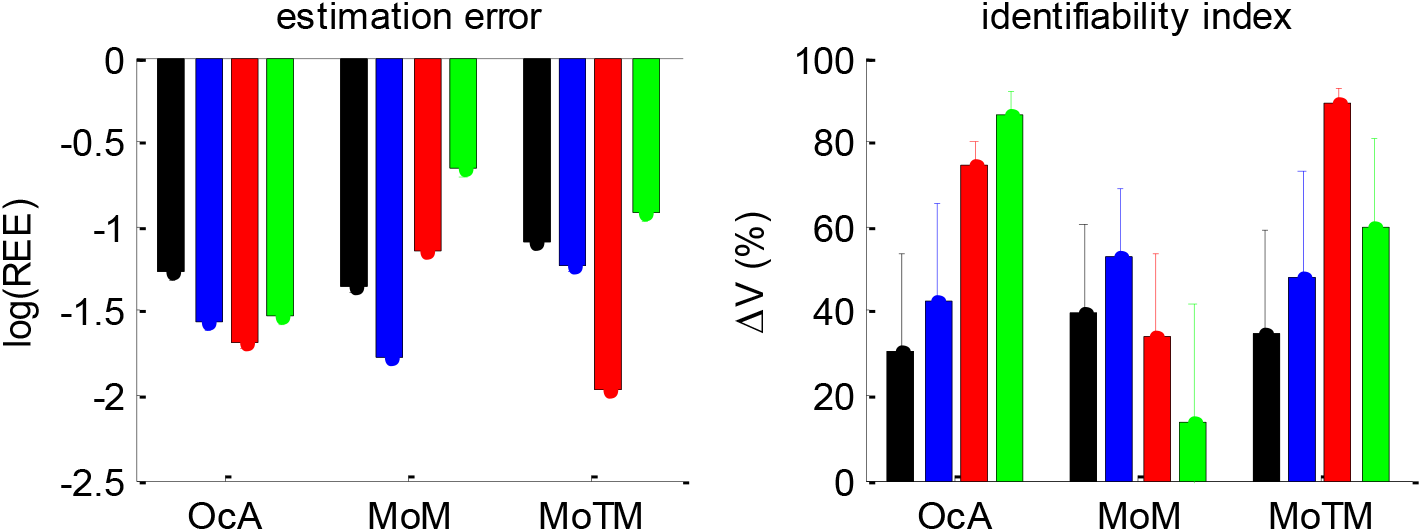
Summary of DDM parameter recovery analyses. **Left panel**: the mean log relative estimation error (y-axis) is shown for all methods (OcA: overcomplete approach, MoM: method of moments, MoTM: method of trial means), and all simulation series (black: full parameter set, blue: fixed drift rate, red: varying drift rates, green: collapsing bounds). **Right panel**: the mean identifiability index Δ*V* (y-axis) is shown for all methods and all simulation series (same format as left panel). Note that the situation in which the full parameter set has to be estimated serves as a reference point. To enable a fair comparison, both the estimation error and the identifiability index are computed for the parameter subset that is common to all simulation series (i.e.: the perturbations ‘standard deviation *σ*, the bound’s height *b*, the initial condition *x*_0_, and the non-decision time *T_ND_*).

Figure 15 enables a visual comparison of the impact of simulation series on parameter estimation methods. As expected, for the method of moments and the method of trial means, the most favourable situation (in terms of estimation error and identifiability) is when the drift rate is fixed and varying over trials, respectively. This is also when these methods perform best in relation to each other. All other situations are detrimental, and eventually yield estimation error and identifiability issues similar or worse than when the full parameter set has to be estimated. This is not the case for the overcomplete approach, which exhibits comparable estimation error and/or identifiability than the best method in all situations, except for collapsing bounds, where it strongly outperforms the two other methods. Here again, we note that parameter recovery for generalized DDMs may, in principle, be improved for the method of moments and/or the method of trial means. But extending these methods to generalized DDMs is beyond the scope of the current work.

## 6. Application to a value-based decision making experiment

To demonstrate the above overcomplete likelihood approach, we apply it to data acquired in the context of a value-based decision making experiment (Lopez-Persem et al., 2016). This experiment was designed to understand how option values are compared when making a choice. In particular, it tested whether agents may have prior preferences that create default policies and shape the neural comparison process.

Prior to the choice session, participants (n = 24) rated the likeability of 432 items belonging to three different domains (food, music, magazines). Each domain included four categories of 36 items. At that time, participants were unaware of these categories. During the choice session, subjects performed series of choices between two items, knowing that one choice in each domain would be randomly selected at the end of the experiment and that they would stay in the lab for another 15 min to enjoy their reward (listening to the selected music, eating the selected food and reading the selected magazine). Trials were blocked in a series of nine choices between items belonging to the same two categories within a same domain. The two categories were announced at the beginning of the block, such that subjects could form a prior or “default” preference (although they were not explicitly asked to do so). We quantified this prior preference as the difference between mean likeability ratings (across all items within each of the two categories). In what follows, we refer to the “default” option as the choice options that belonged to the favored category. Each choice can then be described in terms of choosing between the default and the alternative option.

Figure 16 below summarizes the main effects of a bias toward the default option (i.e. the option belonging to the favored category) in both choice and response time, above and beyond the effect of individual item values.

**Figure 16:**
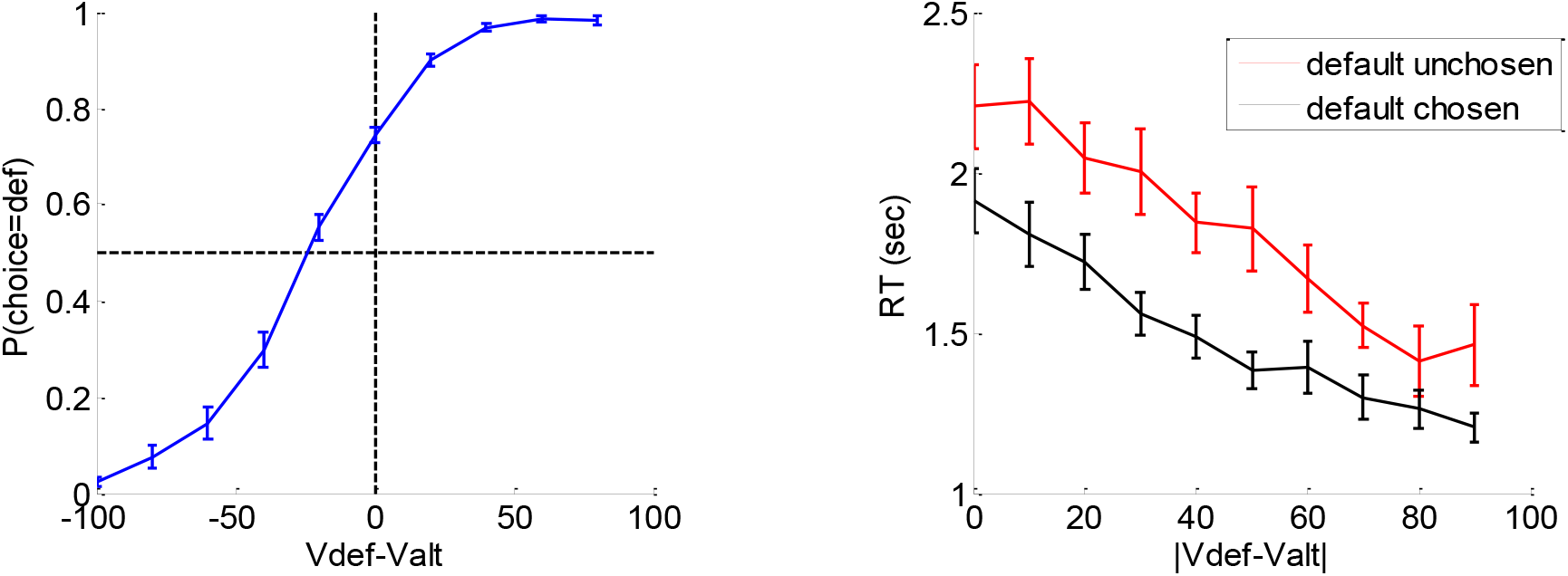
Evidence for choice and RT biases in the default/alternative frame. **Left**: Probability of choosing the default option (y-axis) is plotted as a function of decision value V_def_-V_alt_ (x-axis), divided into 10 bins. Values correspond to likeability ratings given by the subject prior to choice session. For each participant, the choice bias was defined as the difference between chance level (50%) and the observed probability of choosing the default option for a null decision value (i.e. when V_def_=V_alt_). **Right**: Response time RT (y-axis) is plotted as a function of the absolute decision value |V_def_-V_alt_| (x-axis) divided into 10 bins, separately for trials in which the default option was chosen (black) or not (red). For each participant, the RT bias was defined as the difference between the RT intercepts (when V_def_=V_alt_) observed for each choice outcome.

A simple random effect analysis based upon logistic regression shows that the probability of choosing the default option significantly increases with decision value, i.e. the difference V_def_-V_alt_ between the default and alternative option values (t=8.4, dof=23, p<10^−4^). In addition, choice bias is significant at the group-level (t=8.7, dof=23, p<10^−4^). Similarly, RT significantly decreases with absolute decision value |V_def_-V_alt_| (t=8.7, dof=23, p<10^−4^), and RT bias is significant at the group-level (t=7.4, dof=23, p<10^−4^).

To interpret these results, we fitted the DDM using the above overcomplete approach, when encoding the choice either (i) in terms of default versus alternative option (i.e. as is implicit on Figure 10) or (ii) in terms of right option versus left option. In what follows, we refer to the former choice frame as the “default/alternative” frame, and to the latter as the “native” frame. In both cases, the drift rate of each choice trial was set to the corresponding decision value (either V_def_-V_alt_ or V_right_-V_left_). It turns out that within-subject estimates of *σ*, *b* and *T_ND_* do not depend upon the choice frame. More precisely, the cross-subjects correlation of these estimates between the two choice frames is significant in all three cases (*σ*: r=0.76, p<10^−4^; *b*: r=0.82, p<10^−4^; *T*_*ND*_: r=0.94, p<10^−4^). This implies that inter-individual differences in *σ*, *b* and *T_ND_* can be robustly identified, irrespective of the choice frame. However, the between-frame correlation is not significant for the initial bias *x*_0_ (r=0.29, p=0.17). In addition, the initial bias is significant at the group level for the default/alternative frame (F=45.2, dof=[1,23], p<10^−4^) but not for the native frame (F=2.36, dof=[1,23], p=0.14). In brief, the two choice frames only differ in terms of the underlying initial bias, which is only revealed in the default/alternative frame.

Now, we expect, from model simulations, that the presence of an initial bias induces both a choice bias, and a reduction of response times for default choices when compared to alternative choices (cf. upper-left and lower-right panels in Figure 1). The fact that 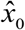 is significant in the default/alternative frame thus explains the observed choice and RT biases shown on Figure 10. But do inter-individual differences in 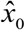 predict inter-individual differences in observed choice and RT biases? The corresponding statistical relationships are summarized on Figure 17 below.

**Figure 17:**
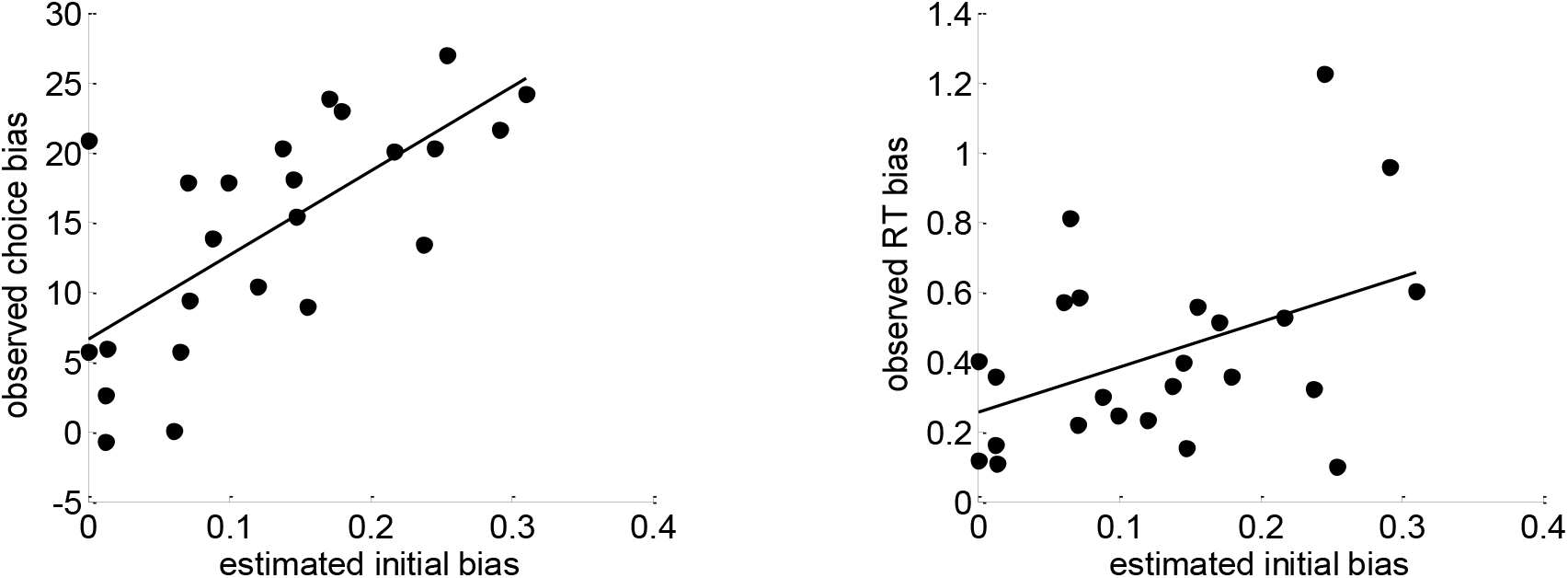
Model-based analyses of choice and RT data. Left: For each participant, the observed choice bias (y-axis) is plotted as a function of the initial bias estimate 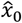 in the default/alternative frame (x-axis). Right: Same for the observed RT bias.

One can see that both pairs of variables are statistically related (choice bias: r=0.70, p<10^−4^; RT bias: r=0.44, p=0.03). This is important, because this provides further evidence in favor of the hypothesis that people's covert decision frame facilitates the default option. Note that this could not be shown using the method of moments or the method of trial means, which were not able to capture these inter-individual differences (see Appendix 7 for details).

Finally, can we exploit model fits to provide a normative argument for why the brain favors a biased choice frame? Recall that, if properly set, the DDM can implement the optimal speed-accuracy tradeoff inherent in making online value-based decisions (Tajima et al., 2016). Here, it may seem that the presence of an initial bias would induce a gain in decision speed that would be overcompensated by the ensuing loss of accuracy. But in fact, the net tradeoff between decision speed and accuracy depends upon how the system sets the bound’s height *b*. This is because *b* determines the demand for evidence before the system commits to a decision. More precisely, the system can favor decision accuracy by increasing *b*, or improve decision speed by decreasing *b*. We thus defined a measure 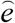 of the optimality of each participant’s decisions, by comparing the speed-accuracy efficiency of her estimated DDM and the maximum speed-accuracy efficiency that can be achieved over alternative bound heights *b* (see Appendix A5 below). This measure of optimality can be obtained either under the default-alternative frame or under the native frame. It turns out that the measured optimality of participants’ decisions is significantly higher under the default/alternative frame than under the native frame (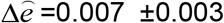, t=2.2, dof=23, p=0.02). In other words, participants’ decisions appear more optimal under the default/alternative frame than under the native frame. We comment on possible interpretations of this result in the Discussion section below.

## 7. Discussion

In this note, we have described an overcomplete approach to fitting the DDM to trial-by-trial RT data. This approach is based upon a self-consistency equation that response times obey under DDM models. It bypasses the computational bottleneck of existing DDM parameter estimation approaches, at the cost of augmenting the model with stochastic neural noise variables that perturb the underlying decision process. This makes it suitable for generalized variants of the DDM, which would not otherwise be considered for behavioral data analysis.

Strictly speaking, the DDM predicts the RT distribution conditional on choice outcomes. This is why variants of the method of moments are not optimal when empirical design parameters (e.g., evidence strength) are varied on a trial-by-trial basis. More precisely, one would need a few trial repetitions of empirical conditions (e.g., at least a few tens of trials per evidence strength) to estimate the underlying DDM parameters from the observed moments of associated RT distributions (Boehm et al., 2018; Ratcliff, 2008; Srivastava et al., 2016). Alternatively, one could rely on variants of the method of trial means to find the DDM parameters that best match expected and observed RTs (Fontanesi et al., 2019a, 2019b; Gluth and Meiran, 2019; Moens and Zenon, 2017; Pedersen et al., 2017; Wabersich and Vandekerckhove, 2014). But this becomes computationally cumbersome when the number of trials is high and one wishes to use generalized variants of the DDM. This however, is not the case for the overcomplete approach. As with the method of trial means, its statistical power is maximal when design parameters are varied on a trial-by-trial basis. But the overcomplete approach does not suffer from the same computational bottleneck. This is because evaluating the underlying self-consistency equation (Equations 7-9) is much simpler than deriving moments of the conditional RT distributions (Broderick et al., 2009; Navarro and Fuss, 2009). In turn, the statistical added-value of the overcomplete approach is probably highest for analysing data acquired with such designs, under generalized DDM variants.

We note that this feature of the overcomplete approach makes it particularly suited for learning experiments, where sequential decisions are based upon beliefs that are updated on a trial-by-trial basis from systematically varying pieces of evidence. In such contexts, existing modelling studies restrict the number of DDM parameters to deal with parameter recovery issues (Frank et al., 2015; Pedersen et al., 2017). This is problematic, since reducing the set of free DDM parameters can lead to systematic interpretation errors. In contrast, it would be trivial to extend the overcomplete approach to learning experiments without having to simplify the parameter space. We will pursue this in forthcoming publications.

Now what are the limitations of the overcomplete approach?

In brief, the overcomplete approach effectively reduces to adjusting DDM parameters such that RT become self-consistent. Interestingly, we derived the self-consistency equation without regard to the subtle dynamical degeneracies that (absorbing) bounds induce on stochastic processes (Broderick et al., 2009). It simply follows from noting that if a decision is triggered at time *τ*, then the underlying stochastic process has reached the bound (i.e. *x_τ_* = ± *b*). This serves to identify the cumulative perturbation that eventually drove the system towards the bound. But a bound hit event at time *τ* is more informative about the history of the stochastic process than just its fate: it also tells us that the path did not cross the barrier before (i.e. |*x_t_*| < *b* ∀*t* < *τ*). This disqualifies those sample paths whose first-passage time happens sooner, even though all barrier crossings are (by definition) “self-consistent”. In retrospect, one may thus wonder whether the self-consistency equation may be suboptimal, in the sense of incurring some loss of information. Critically however, no information is lost about cumulative perturbations (or about DDM parameters). Although these are not sufficient to discriminate between the many sample paths that are compatible with a given RT, this is essentially irrelevant to the objective of the overcomplete approach. In turn, the existing limitations of the overcomplete approach lie elsewhere.

First and foremost, the self-consistency equation cannot be used to simulate data (recall that RTs appear on both the left- and right-hand sides of the equation). This restricts the utility of the approach to data analysis. Note however, that data simulations can still be performed using Equation 2, once the model parameters have been identified. This enables all forms of posterior predictive checks and/or other types of model fit diagnostics (Palminteri et al., 2017). Second, the accuracy of the method depends upon the reliability of response time data. In particular, the recovery of the noise's standard deviation depends upon the accuracy of the empirical proxy for decision times (cf. second term in Equation 7). Third, the computational cost of model inversion scales with the number of trials. This is because each trial has its own nuisance perturbation parameter. Note however, that the ensuing computational cost is many orders of magnitude lower than that of standard methods for generalized DDM variants. Fourth, proper bayesian model comparison may be more difficult. In particular, simulations show that a chance model always has a higher model evidence than the overcomplete model. This is another consequence of the overcompleteness of the likelihood function, which eventually pays a high complexity penalty cost in the context of Bayesian model comparison. Whether different DDM variants can be discriminated using the overcomplete approach is beyond the scope of the current work.

Let us now discuss the results of our model-based data analysis from the value-based decision making experiment (Lopez-Persem et al., 2016). Recall that we eventually provided evidence that peoples’ decisions are more optimal under the default/alternative frame than under the native frame. Recall that this efficiency gain is inherited from the initial condition parameter *x*_0_, which turns out be significant under the default/alternative frame. The implicit interpretation here is that the brain’s decision system starts with a prior bias towards the default option. Critically however, we would have obtained the exact same results, would we have fixed the initial condition to zero but allowed upper and lower decision bounds to be asymmetrical. This is interesting, because it highlights a slightly different interpretation of our results. Under this alternative scenario, one would state that the brain’s decision system is comparatively less demanding regarding the evidence that is required for committing to the default option. In turn, the benefit of lowering the bound for the default option may simply be to speed up decisions when evidence is congruent with default preferences, at the expense of slowing down incongruent decisions. Importantly, this strategy does not compromise decision accuracy if the incongruent decisions are rarer than the congruent ones (as is effectively the case in this experiment).

At this point, we would like to discuss potential neuroscientific applications of trial-by-trial estimates of “neural” perturbation terms. Recall that the self-consistency equation makes it possible to infer these neural noise variables from response times (cf. Equations 7 or 9). For the purpose of behavioral data analysis, where one is mostly interested in native DDM parameters, these are treated as nuisance variables. However, should one acquire neuroimaging data concurrently with behavioral data, one may want to exploit this unique feature of the overcomplete approach. In brief, estimates of “neural” perturbation terms moves the DDM one step closer to neural data. This is because DDM-based analysis of behavioral data now provides quantitative trial-by-trial predictions of an underlying neural variable. This becomes particularly interesting when internal variables (e.g., drift rates) are systematically varied over trials, hence de-correlating the neural predictor from response times. For example, in the context of fMRI investigations of value-based decisions, one may search for brain regions whose activity eventually perturbs the computation and/or comparison of options’ values. This would extend the portfolio of recent empirical studies of neural noise perturbations to learning-relevant computations (Drugowitsch et al., 2016; Findling et al., 2019; Wyart and Koechlin, 2016). Reciprocally, using some variant of mediation analysis (Brochard and Daunizeau, 2020; Lindquist, 2012; MacKinnon et al., 2007), one may extract neuroimaging estimates of neural noise that can inform DDM-based behavioral data analysis. Alternatively, one may model neural and behavioral data in a joint and symmetrical manner, with the purpose of testing some predefined DDM variant (Rigoux and Daunizeau, 2015; Turner et al., 2015).

Finally, one may ask how generalizable the overcomplete approach is? Strictly speaking, one can evaluate the self-consistency equation under any DDM variant, as long as the mapping *z*: *x* ⟶ *z* (*x*) from the base random walk to the bound subspace is invertible (cf. Equations 8-9). No such formal constraint exists for the dynamical form of the collapsing bound. This spans a family of DDM variants that is much broader than what is currently being used in the field (Fengler et al., 2020; Shinn et al., 2020). For example, this family includes decision models that trigger a decision when decision *confidence* reaches a bound (Lee and Daunizeau, 2020; Tajima et al., 2016). To the best of our knowledge, there is not a single example of existing DDM variants that does not belong to this class. Having said this, future extensions of the DDM framework may render the current overcomplete approach obsolete. Our guess is that such DDM improvements may then need to be informed with additional behavioral data, such as decision confidence (De Martino et al., 2012) and/or mental effort (Lee and Daunizeau, 2020), for which other kinds of self-consistency equations may be derived.

To conclude, we note that the code that is required to perform a DDM-based data analysis under the overcomplete approach will be made available soon from the VBA academic freeware https://mbb-team.github.io/VBA-toolbox/ (Daunizeau et al., 2014).

## Acknowledgements

We would like to thank Dr. Alizée Lopez-Persem for providing us with the empirical data that serves to demonstrate our approach.

## Appendix 1 Summary of the variational Laplace approach

To begin with, note that the mathematical form of all generative models *m* that we consider here can be written as follows (for notational convenience, we ignore the dependence on decision outcomes):

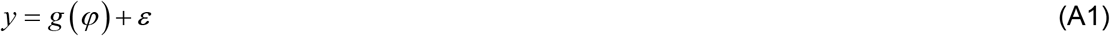

where *y* are the fitted data (typically: RTs or RT moments), *φ* are model parameters, *ε* are model residuals, and *g* is a mapping that depends upon the generative model. For the overcomplete approach, the observation mapping *g* is given by the self-consistency equation, having replaced native DDM parameters with their corresponding dummy variables *φ* (cf. sections 3.a and 3.b of the main text). The observation mapping of other approaches are described in Appendix 2 and 3 below.

Recall that the variational Laplace scheme is an iterative algorithm that indirectly optimizes an approximation to both the model evidence *p* (*y | m*) and the posterior density *p* (*φ* | *y*, *m*).

The key trick is to decompose the log model evidence into:

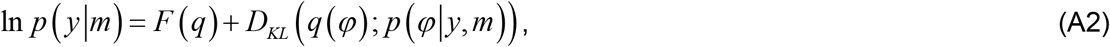

where *q*(*φ*)is any arbitrary density over the model parameters, *D_KL_* is the Kullback-Leibler divergence and the so-called *free energy F*(*q*) is defined as:

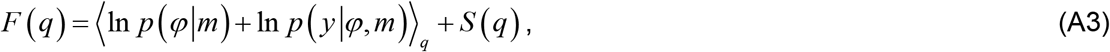

where *S* (*q*) is the Shannon entropy of *q* and the expectation 〈•〉_*q*_ is taken under *q*.

From equation A2, maximizing the functional *F* (*q*) w.r.t. *q* indirectly minimizes the Kullback-Leibler divergence between *q* (*φ*) and the exact posterior *p* (*φ*|*y*, *m*). This decomposition is complete in the sense that if *q* (*φ*) = *p* (*φ*| *y*, *m*), then *F* (*q*) = ln *p* (*y|m*).

The variational Laplace algorithm iteratively maximizes the free energy *F* (*q*) under simplifying assumptions (see below) about the functional form of *q*, rendering *q* an approximate posterior density over model parameters and *F* (*q*) an approximate log model evidence (Daunizeau, 2017; Friston et al., 2007). The free energy optimization is then made with respect to the sufficient statistics of *q*, which makes the algortithm generic, quick and efficient.

Under normal i.i.d. model residuals, the likelihood function writes:

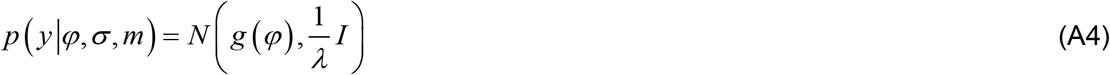

where *λ* is the residuals’ precision or inverse variance hyperparameter.

We also use Gaussian priors *p* (*φ* | m) = *N* (*μ*_0_, ∑_0_) for model parameters and gamma priors for precision hyperparameters *p* (*λ* | *m*) = *Ga* (*ϖ*_0_, *κ*_0_)

In what follows, we derive the variational Laplace algorithm under a “mean-field” separability assumption between parameters and hyperparameters, i.e.: *q* (*φ*, *λ*) = *q* (*φ*) *q* (*λ*). We will see that this eventually yields a Gaussian posterior density *q* (*φ*) ≈ *N* (*μ*, ∑) on model parameters, and a Gamma posterior density *q* (*λ*) = *Ga* (*ϖ, κ*) on the precision hyperparameter.

First, let us note that, under the Laplace approximation, the free energy bound on the log-model evidence can be written as:

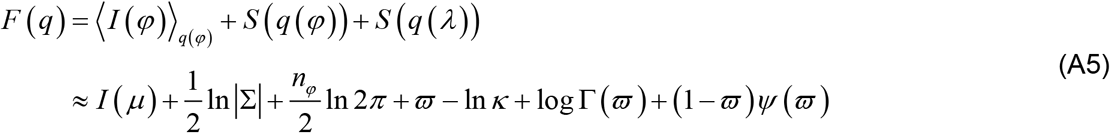

where *n_φ_* is the number of parameters, Γ(•) is the gamma function, *ψ*(•) is the digamma function, and *I* (*φ*) is defined as:

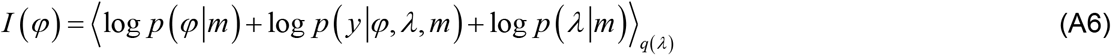

Given the Gamma posterior *q* (*λ*) on the precision hyperparameter, *I* (*φ*) can be simply expessed as follows:

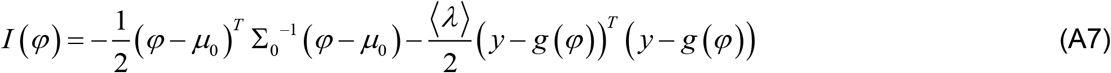

where we have ignored the terms that do not depend upon *φ*, and 〈*λ*〉 = *E*[*λ* | *y, m*] = *ϖ/κ* is the posterior mean of the data precision hyperparameter *λ*.

The variational Laplace update rule for the approximate posterior density *q* (*φ*) on model parameters now simply reduces to an update rule for its sufficient statistics:

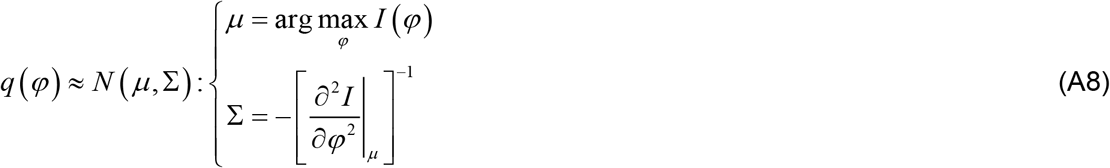

In Equation A8, the first-order moment *μ* of *q* (*φ*) is obtained from the following Gauss-Newton iterative gradient ascent scheme:

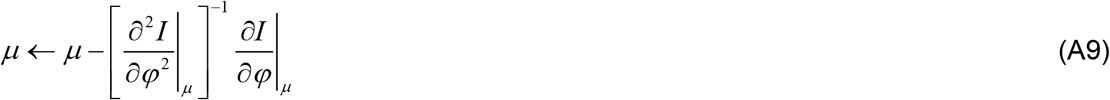

where the gradient and Hessians of *I* (*φ*)are given by:

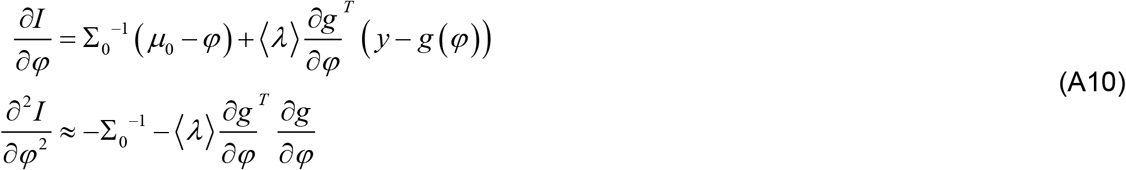

At convergence of the above gradient ascent, the approximate posterior density *q* (*φ*) on the precision hyperparameter is updated as follows:

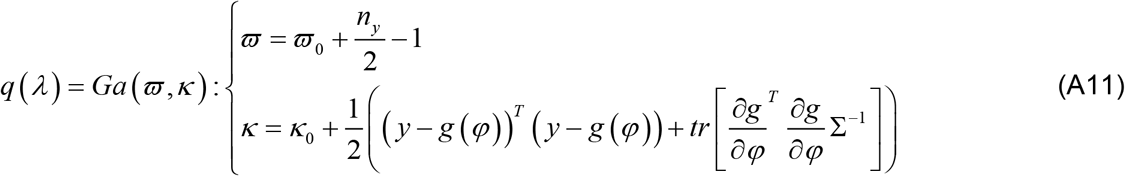

where *n_y_* is the number of data samples.

The variational Laplace scheme alternates between Equations A8 and A11 iteratively until convergence of the free energy.

At this point, we note that, for all estimation methods, we set the prior probability density functions as follows:

- *p* (*φ*_*i*_ | *m*) = *N* (0,1) ∀*i*, i.e. the prior mean of model parameters is *μ*_0_ = 0 and their prior variance is ∑_0_ = *I*. For group studies, this prior setting can be replaced with parametric empirical Bayes priors (Daunizeau, 2019).
- *p* (*λ* | *m*) = *Ga* (1, *κ*_0_), where *κ*_0_ = *V* [*y*] is the variance of the dependent variable *y*. In other words, the data is *a priori* assumed to be driven entirely by noise. This ensures that the prior and likelihood components of *I* (*φ*) are balanced when the variational Laplace algorithm starts.

This completes the description of the variational Laplace approach.

For more details, we refer the interested reader to the existing literature on variational approaches to approximate bayesian inference (Beal, 2003; Daunizeau, 2017; Friston et al., 2007). We note that the above variational Laplace approach is implemented in the opensource VBA toolbox (Daunizeau et al., 2014).

## Appendix 2 Method of moments

Let *p* (*τ* | *o*, *v*, *x*_0_, *b*, *σ*) be the theoretical probability density function of hitting times *τ*, conditional on the decision outcome *o* and DDM parameters. Here, we evaluate *p* (*τ* | *o*, *v*, *x*_0_, *b*, *σ*) using a fast and accurate semi-analytical approach which was derived for the “vanilla” DDM case (Navarro and Fuss, 2009). Although not generalizable to more complex DDM variants, this eschews the need for sampling scheme or numerical solvers of the underlying Fokker-Planck equation, which are computationally very demanding.

Then, RT conditional (central) moments 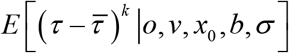 are simply given by:

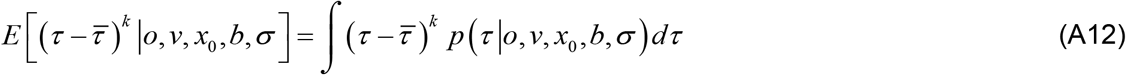

where *k* is the moment’s order (e.g., *k* = 2: variance, *k* = 3: skewness), 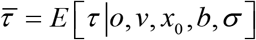 is the conditional mean response time. We evaluate the right-hand term of Equation A12 using numerical integration, on a predefined integration grid.

The method of moments then attempts to match the empirical and theoretical moments of RT data (conditional on decision outcomes). This is formalized through a generative model of the form given in Equation A1 above, where *y* is the set of empirical RT moments (here, up to order 3) and the observation mapping *g* (•) is made of the corresponding set of theoretical moments (cf. Equation A12). The variational Laplace approach then yields an approximate posterior density on DDM parameters, under the assumptions given in Appendix 1. This completes the summary description of the method of moments.

The method of moments is known to be fast and robust, which is why it is an established approach for inference on population parameters in the statistical literature (Newey and West, 1987). In the context of DDM applications (Grasman et al., 2009; Pedersen and Frank, 2020; Vandekerckhove and Tuerlinckx, 2008; Voss and Voss, 2007; Wagenmakers et al., 2007, 2008; Wiecki et al., 2013), it lumps all trial-by-trial variations into statistical moments of the RT distribution. Importantly, this may induce some severe information loss in some contexts.

For example, let us consider experimental designs where DDM drift rates are varied systematically over trials. The method of moments would account for such variations by marginalizing the conditional RT distribution over the drift rate trial-by-trial distribution:

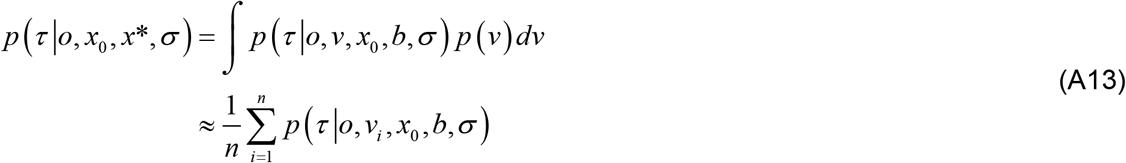

where *n* is the number of trials, *v_i_* is the predicted drift rate at trial *i*, and *p* (*τ* | *o*, *v_i_*, *x*_0_, *b*, *σ*) is the corresponding RT distribution. One can see that the resulting marginal distribution *p* (*τ* | *o*, *x*_0_, *b*, *σ*) will be blind to many forms of predictable trial-by-trial drift rate variations. To begin with, *p* (*τ* | *o*, *x*_0_, *b*, *σ*) is invariant to any arbitrary permutation of the trials. In addition, as we will see below, increasing drift rate variability has almost the same effect on the marginal distribution *p* (*τ* | *o*, *x*_0_, *b*, *σ*) than increasing the variance *σ*_2_ of neural stochastic perturbation terms. This is summarized on Figure A1 below.

**Figure A1:**
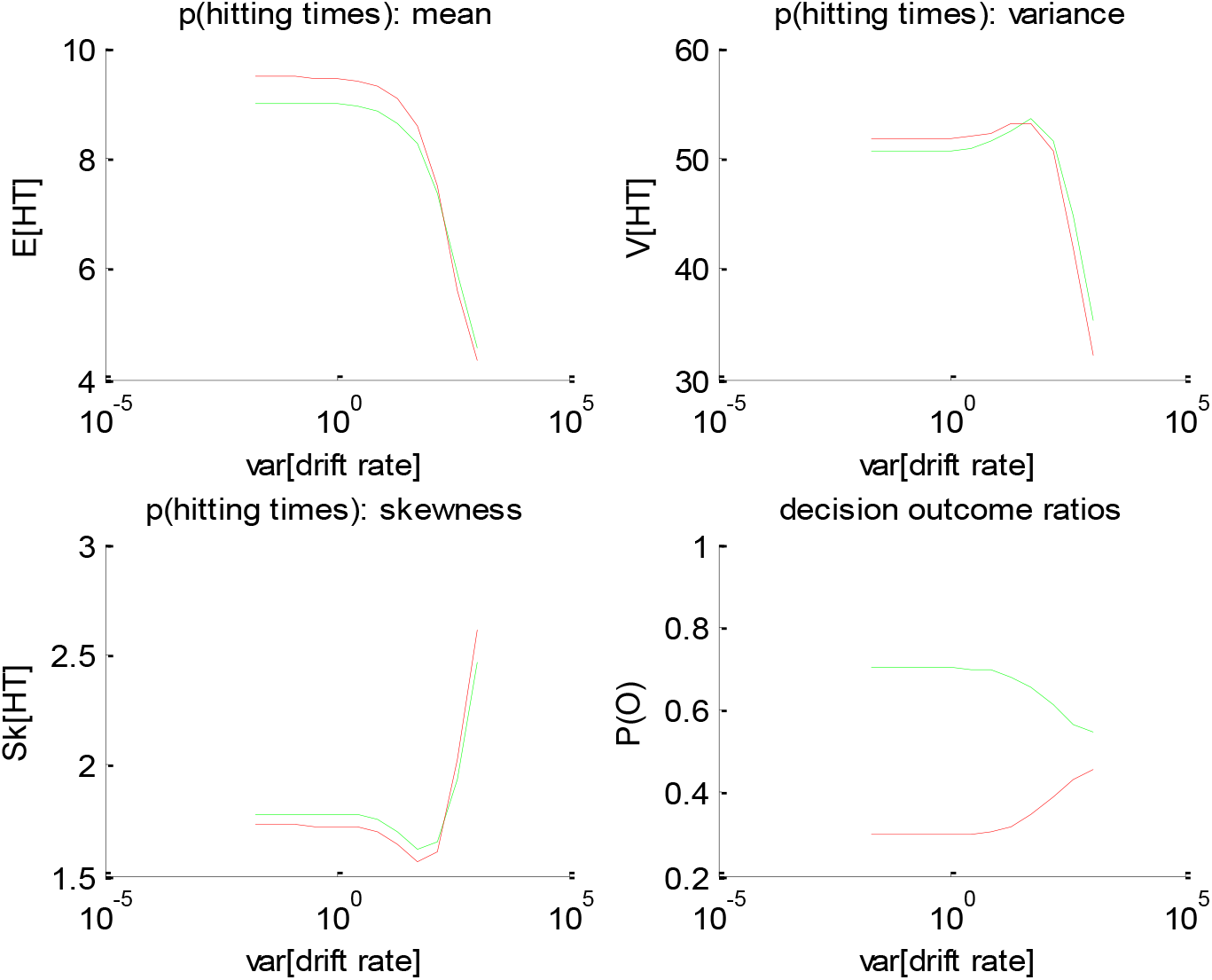
impact of the trial-by-trial variance of drift rate. Same format as Fig 1 of the main text. Note that the mean drift rate 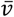 was fixed to its default value, i.e.: 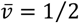, which corresponds to black dotted lines on all panels of Figures 1 to 4).

As one can see, the trial-by-trial variance of drift rate essentially has no impact on the marginal distribution *p* (*τ* | *o*, *x*_0_, *b*, *σ*) as long as it is below or of the same order of magnitude than the mean drift rate. From this point onwards, it decreases the hitting time's mean and its variance, increases its skewness, and increases the entropy of the outcome probability. This effect is qualitatively identical to the impact of the noise’s standard deviation *σ* (cf. Figure 3 of the main text). Practically speaking, Figure A1 implies that the method of moments may confuse trial-by-trial variations of drift rates with the effect of stochastic perturbations. In retrospect, this is expected, because any form of trial-by-trial drift-rate variations will eventually conspire with stochastic perturbations to inflate trial-by-trial RT variations. We note that this argument generalizes to trial-by-trial variations in other model parameters.

## Appendix 3 Method of the trial means

A possible improvement over the above method of moments is to match trial-by-trial RT data to their corresponding theoretical mean. This is typically done by evaluating trial-specific likelihood functions from samplers of the DDM model or related numerical approaches, which may then be used within a probabilistic framework for parameter estimation (Fontanesi et al., 2019a, 2019b; Gluth and Meiran, 2019; Moens and Zenon, 2017; Pedersen et al., 2017; Wabersich and Vandekerckhove, 2014). For numerical expediency, we here rather rely on the variational Laplace described in Appendix 1. In brief, the method of trial means relies upon a generative model of the form given in Equation A1, where *y* is now the raw RT data trial series and the observation mapping *g* (•) is made of the corresponding set of trial-by-trial expected RTs (conditional on decision outcomes). The latter are derived from the following analytical expression for the RT conditional mean, which is valid under the “vanilla” DDM (Srivastava et al., 2016):

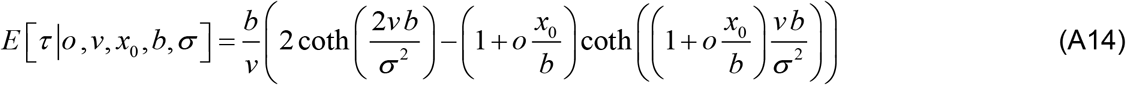

where *o* ∈ {−1, 1} is the decision outcome. One can then evaluate Equation A14 for each trial, given its corresponding set of DDM parameters. In particular, if one knows how drift rates vary over trials, then one can predict the ensuing expected RT variations. The variational Laplace treatment of the ensuing generative model then yields an approximate posterior density on the remaining DDM parameters. This completes the summary description of the method of trial means.

## Appendix 4 parameter recovery analysis

Parameter recovery analyses aim at evaluating how accurate parameter estimates are. This is done by comparing simulated and estimated parameters using Monte-Carlo simulation series, which we now detail.

Let *θ* = {*v*, *x*_0_, *b*, *σ*,*T_ND_*} be the set of all DDM parameters. In what follows, we refer to *θ*_ij_ *i*^th^ element of *θ* at the *j*^th^ Monte-Carlo simulation. At each Monte-Carlo simulation, the vector *θ_•j_* 0f simulated parameters is randomly drawn from a standard normal distribution (passed through its corresponding mapping, see section 3.b of the main text). Given *θ_•j_*, we then simulate a series of N=200 DDM trials according to Equation 2. The ensuing trial series of choice outcomes and response times is then fed to each parameter estimation method, each of which outputs a vector 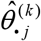 of estimated parameters, where *k* is the estimation method’s index. We repeat this procedure *N_mc_* = 1000 times, yielding a series of 1000 simulated parameter sets, and their corresponding 1000 estimated parameters sets (per estimation method). Should 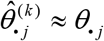, then parameter recovery would be perfect. The accuracy of parameter recovery can thus be visually evaluated when plotting estimated parameter against simulated parameters (cf. Figures 7, 9, 11 and13).

To further quantifying the accuracy of parameter recovery, we measure the mean relative estimation error 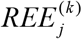 for each Monte-Carlo simulation and each estimation method:

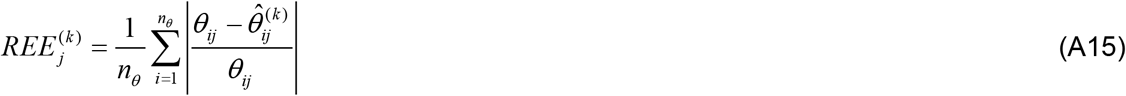

where *n_θ_* is the number of DDM parameters to be estimated (see main text). Between-method comparisons can then be compared using pairwise statistical tests. We then use one-sample F-tests on the difference 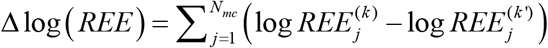 of log-transformed estimation errors. Log-transformed estimation errors are summarized on Figure 15 of the main text.

We also quantify pairwise non-identifiability issues, which arise when the estimation method confuses two parameters with each other. We do this using so-called “recovery matrices”, which summarize whether variations (across the 1000 Monte-Carlo simulations) in estimated parameters faithfully capture variations in simulated parameters. We first z-score simulated and estimated parameters across Monte-Carlo simulations. We then regress each estimated parameter against all simulated parameters through the following multiple linear regression model:

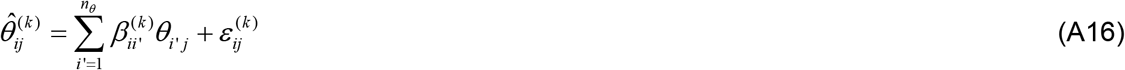

where 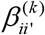 are regression weights, and 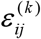 are regression residuals. Here, regression weights are partial correlation coefficients between simulated and estimated parameters (across Monte-Carlo simulations). More precisely, 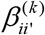 quantifies the impact that variations of the simulated parameter *θ*_i’•_ have on variations of the estimated parameter 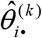 conditional on all other simulated parameters. Would parameters be perfectly identifiable, then 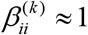 and 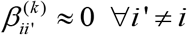. Recovery matrices simply report the squared regression weights 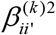 for each simulated parameter (cf. Figure 8, 10, 12 and 14 of the main text).

Note that the regression model in Equation A16 effectively decomposes the observed variability in the estimated parameter 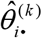 into “correct variations” that are induced by variations in the corresponding simulated parameter *θ*_i•_, and “incorrect variations” that are induced by the remaining simulated parameters *θ*_i’•_ (with *i*’ ≠ *i*). In turn, this enables the derivation of the following identifiability index Δ*V*_(*k*)_:

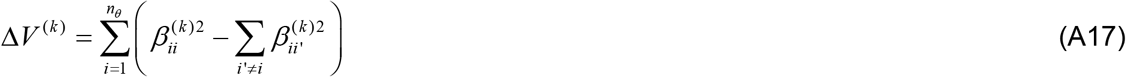

where the first and the second summands of the right-hand term of Equation A17 are the amount of “correct” and “incorrect” variations, respectively. The identifiability index is maximal when the amount of “correct variations” dominates the amount of “incorrect variations”. Mean identifiability indices are summarized on Figure 15 of the main text.

## Appendix 5 evaluating the computational optimality of a DDM

Recall that the DDM has been shown to be the optimal solution to the computational problem of trading speed with accuracy when making online value-based decisions (Tajima et al., 2016). This speed-accuracy tradeoff arises because the decision system needs to accumulate noisy value signals before making a reliable decision. Arguably, the decision system cannot control the upstream decision variables, such as the evidence strength (i.e. the drift rate *v*), or the reliability of value signals (i.e. the noise’s standard deviation *σ*). However, it can set the bound’s height *b* so as to optimize the ensuing speed-accuracy tradeoff. More precisely, the system can favor decision accuracy by increasing *b*, or improve decision speed by decreasing *b*. In context of perceptual decision making, one can score the efficiency of any *b* setting, in terms of the ensuing reward rate *RR* (*b*) (Balci et al., 2011; Bogacz et al., 2006; Zhang, 2012):

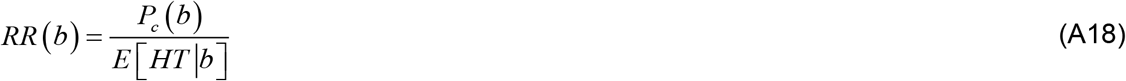

where *E* [*HT | b*] is the expected hitting time, 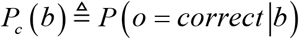 is the ensuing probability of making a correct decision, and we have neglected non-decision times and inter-trial intervals. The “optimal” bound’s height is such that it maximizes the reward rate. Now the system may not have set the bound’s height to its optimal value. The optimality 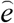 of the system can thus be measured in terms of the ratio between the actual efficiency of the system and its optimal efficiency:

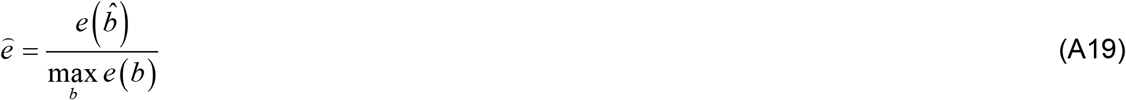

where 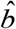 is the estimated bound’s height of a given participant. Equation A19 can be extended to situations where drift rate changes in a trial-by-trial fashion, by averaging the optimality score 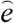 over trials. This makes the average optimality score 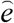 a measure of how well the system adapts to the global statistics of decisions it has to make.

Figure A2 below shows a typical example of the derivation of the optimality 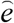 score, based upon the estimated DDM parameters of a study participant (and the corresponding sequence of drift rates over trials) under the default/alternative frame of reference.

**Figure A2:**
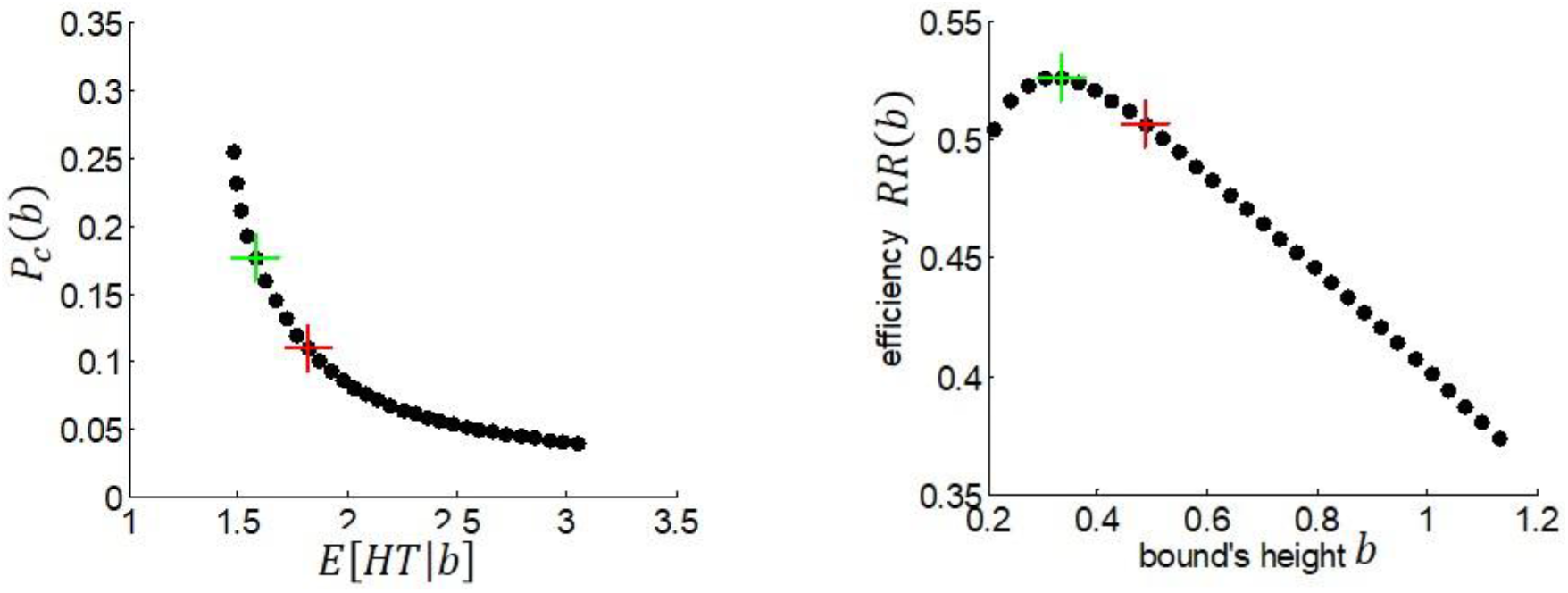
Model-based evaluation of DDM optimality. **Left panel**: The probability *P*_*c*_(*b*) of making a correct decision (y-axis) is plotted against the expected decision time *E*[*HT | b*] (y-axis) for each possible bound height. The green and red crosses depict the optimal and actual speed-accuracy tradeoffs, respectively. **Right panel**: The efficiency *e*(*b*)(y-axis) is plotted for each possible bound height *b*. The green cross depicts the optimal efficiency and the red cross depicts the actual efficiency achieved for this participant.

The optimality score of this participant is estimated as 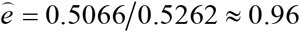. In other terms, this participant’s decisions seem to arise from a DDM decision system that is close to its optimal setting, in terms of the underlying speed-accuracy tradeoff.

## Appendix 6 The “ballistic” limit of drift rates

In this work, we assert that drift rates are bounded above and below (cf. section 3.b if the main text). More precisely, we enforce the following constraint on drift rate estimators:

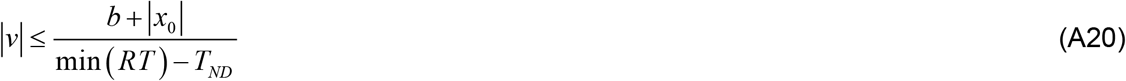

where the minimum RT is taken over trials. In what follows, we refer to the right-hand term of Equation A20 as the “ballistic” limit of drift rates.

So-called “linear ballistic accumulation models” can be understood as non-stochastic variants of the DDM, whereby evidence accumulation follows a straight line with a slope equal to the DDM's drift rate (Goldfarb et al., 2014; Osth et al., 2017). Under such ballistic models, it is not unreasonable to interpret RT variability in terms of drift rate variability. In turn, drift rates would necessarily obey Equation A20.

Intuitively, this result generalizes to DDMs because the net effect of adding stochastic perturbations to ballistic evidence accumulation is to accelerate decision times. Consider the vanilla DDM, where the drift rate is fixed but unknown. If there is no decision error, then the RT distribution is approximately centred around the corresponding “ballistic” RT. In turn, a “ballistic” drift rate estimate based upon the minimum RT would necessarily be greater than the true drift rate, hence the “ballistic limit”. This reasoning is also valid for decision errors, though there is no trivial ballistic equivalent to decision errors.

To check this reasoning, we performed a series of 2000 Monte-Carlo simulations, where we varied randomly all DDM parameters (without imposing the “ballistic limit” to drift rates). Figure A3 below compares the (unconstrained, randomly drawn) drift rates to their theoretical “ballistic limit”.

**Figure A3:**
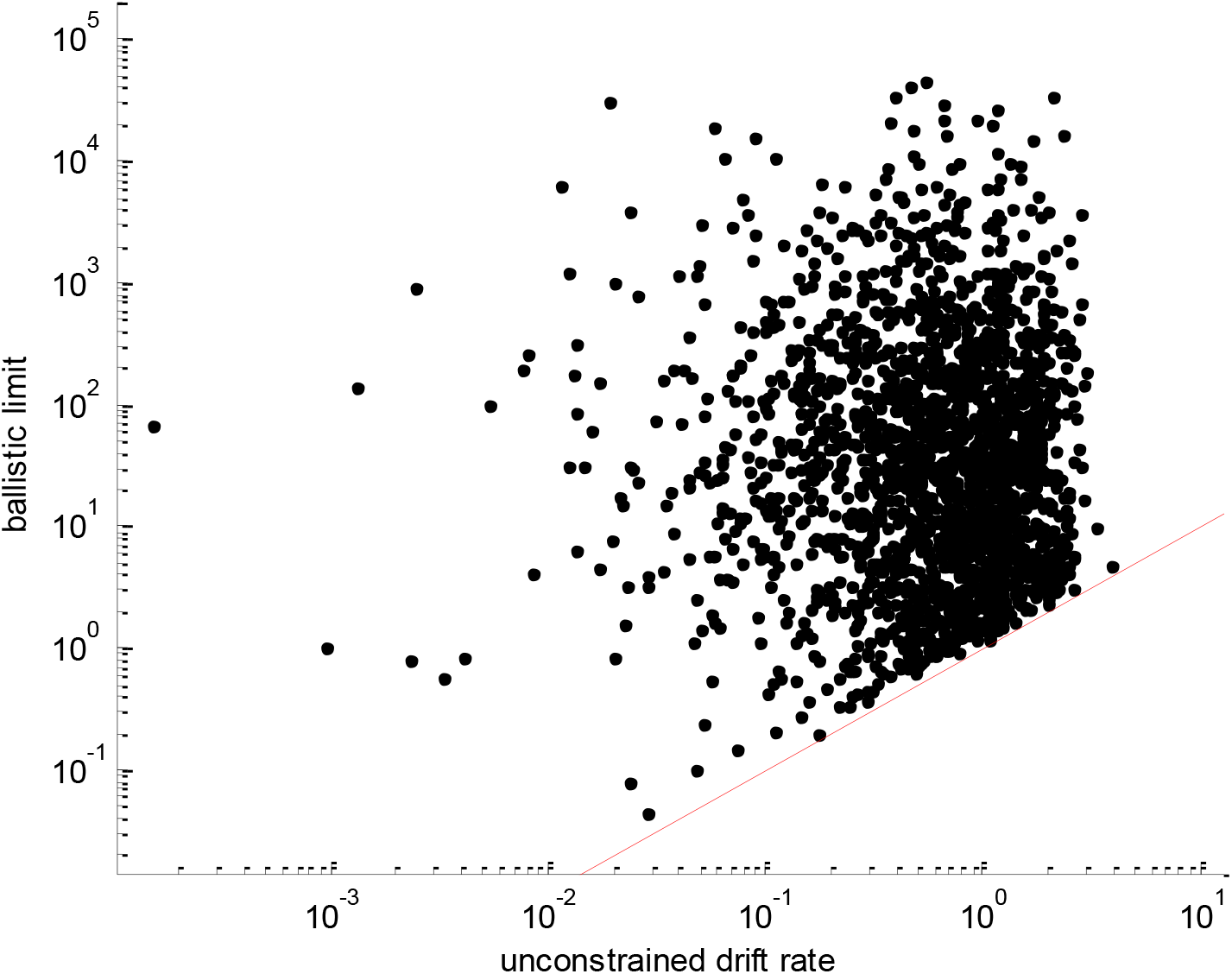
Validity of the “ballistic limit” of drift rates. The right-hand side of Equation A20 (ballistic limit, y-axis) is plotted against the left-hand-side of Equation A20 (absolute value of unconstrained drift rate, x-axis). Each dot is a Monte-Carlo simulation of 200 trials of a DDM with randomly drawn parameters. The red line indices the identity mapping, i.e. equality between drift rates and their ballistic limit.

One can see that the ballistic limit is always greater than unconstrained drift rates, irrespective of the actual DDM parameter setting. This validates the “ballistic” limit of drift rates.

## Appendix 7 Prior preferences: alternative analysis results

In the main text, we report the results of value-based decision data analysis using the overcomplete approach. Here, we summarize the results of analyses of the same dataset, this time using the method of moments and the method of trial means. Note that we used the exact same statistical testing procedures in all cases.

To begin with, we tested the group-level significance of estimated initial conditions 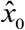. For all estimation methods, it is significant for the default frame (method of moments: F=386.2, dof=[1,23], p<10^−4^, method of trial means: F=60.1, dof=[1,23], p<10^−4^) but not for the native frame (method of moments: F=0.81, dof=[1,23], p=0.38, method of trial means: F=1.82, dof=[1,23], p=0.19). This result is qualitatively similar to the overcomplete approach.

We then asked whether inter-individual differences in 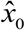 predict inter-individual differences in observed choice and RT biases. Figure A4 and A5 below summarize these relationships for method of moments and the method of trial means, respectively.

**Figure A4:**
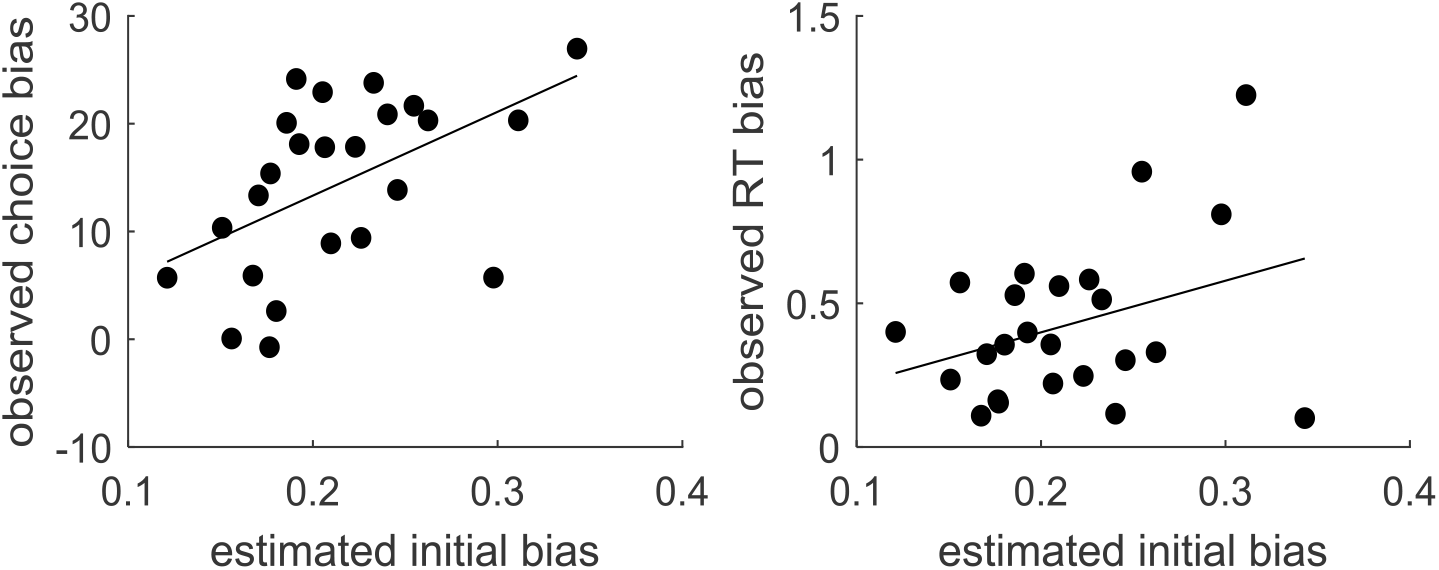
Model-based analyses of choice and RT data: method of moments. Same format as Figure 17.

**Figure A5:**
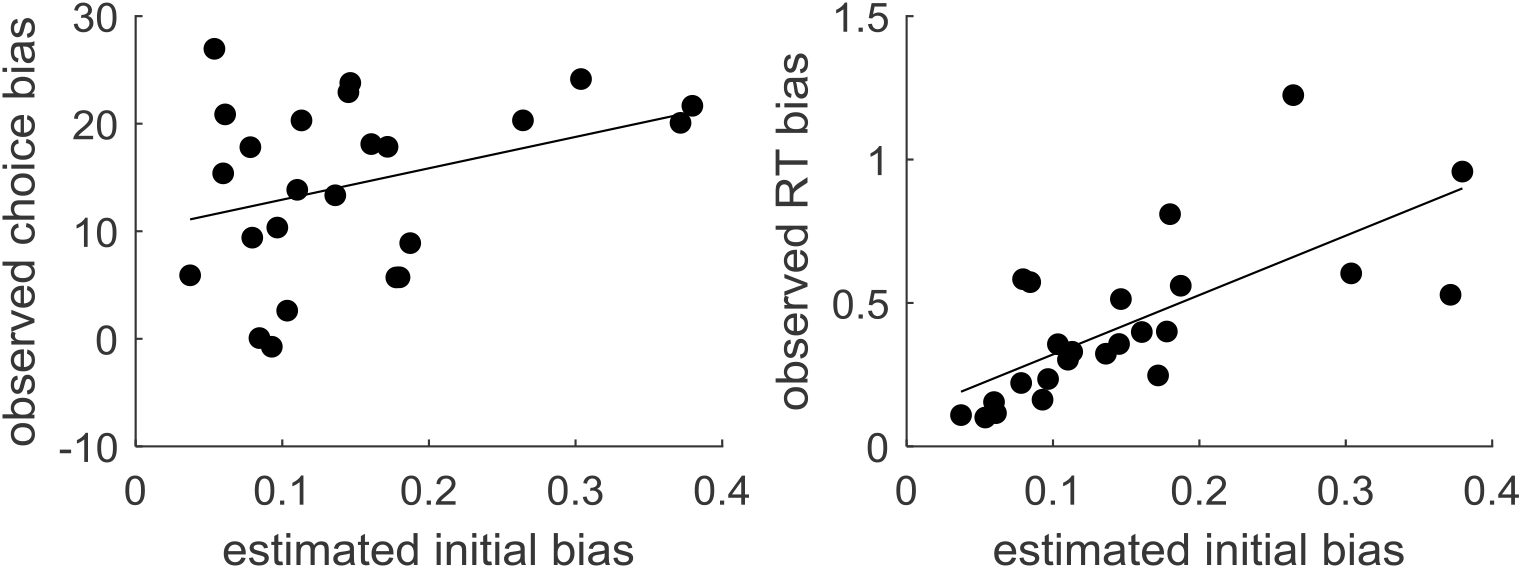
Model-based analyses of choice and RT data: method of trial means. Same format as Figure 17.

When estimated with the method of moments, inter-individual differences in 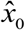 significantly correlates with choice bias (r=0.51, p=0.01) but not with RT bias (r=0.34, p=0.11).

When estimated with the method of trial means, inter-individual differences in 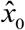 significantly correlates with RT bias (r=0.70, p=0.0002) but not with choice bias (r=0.34, p=0.11).

Taken together, these results imply that the overcomplete approach provides estimates of inter-individual differences that more reliably relate to observable biases than the method of moments or the method of the trial means.

Lastly, we note that that, using the overcomplete approach, analyzing the data under a generalized variant of the DDM (with, e.g., collapsing bounds) yields qualitatively similar results as under the vanilla DDM. However, the bound’s exponential decay rate was not significant at the group-level (F=0.89, dof=[1,23], p=0.35). In other terms, there is no strong evidence for exponentially decaying bounds in this dataset, which is why we report the results of the vanilla DDM in the main text.

